# Critical re-evaluation of the slope factor of the operational model of agonism: Do not exponentiate operational efficacy

**DOI:** 10.1101/2021.04.07.438803

**Authors:** Jan Jakubík

## Abstract

Although being a relative term, agonist efficacy is a cornerstone in the proper assessment of agonist selectivity and signalling bias. The operational model of agonism (OMA) has become successful in the determination of agonist efficacies and ranking them. In 1983, Black and Leff introduced the slope factor to the OMA to make it more flexible and allow for fitting steep as well as flat concentration-response curves. Functional analysis of OMA demonstrates that the slope factor implemented by Black and Leff affects relationships among parameters of the OMA. Fitting of the OMA with Black & Leff slope factor to concentration-response curves theoretical model-based data resulted in wrong estimates of operational efficacy and affinity. In contrast, fitting the OMA modified by the Hill coefficient to the same data resulted in correct estimates of operational efficacy and affinity. Therefore, OMA modified by the Hill coefficient should be preferred over the Black & Leff equation for ranking of agonism and subsequent analysis, like quantification of signalling bias, when concentration-response curves differ in the slope factor and mechanism of action is known. Otherwise. Black & Leff equation should be used with extreme caution acknowledging potential pitfalls.

## Introduction

Since its introduction in 1983, the operational model of agonism (OMA) has become a golden standard in the evaluation of agonism and subsequently also of signalling bias(Black and Leff, 1983). The OMA describes the response of the system as a function of ligand concentration using three parameters: 1) The equilibrium dissociation constant of agonist (K_A_) to the receptor initiating functional response; 2) The maximal possible response of the system (E_MAX_); 3) The operational efficacy of agonist (τ). Original OMA describes the dependence of functional response on the concentration of ligand by rectangular hyperbola (Eq. 2). In practice, positive cooperativity or positive feedback lead to steep and negative cooperativity or negative feedback lead to flat concentration-response curves that the OMA does not fit to. Therefore, an equation intended for the description of non-hyperbolic concentration curves (Eq. 7), particularly for systems where the number of receptors varies, was introduced by Black et al.(Black et al., 1985). Since then, Eq. 7 is commonly used. The presented mathematical analysis of Eq. 7 shows that the slope factor affects the relationship between observed maximal response to agonist and operational efficacy and the relationship between the concentration of agonist for half-maximal response and its equilibrium dissociation constant.

The Hill–Langmuir equation was originally formulated to describe the cooperative binding of oxygen molecules to haemoglobin(Gesztelyi et al., 2012). The Hill coefficient reflects the sense and magnitude of cooperativity among concurrently binding ligands. A value of the Hill coefficient lower than 1 indicates negative cooperativity that is manifested as a flat concentration-response curve. The smaller value of the Hill coefficient the flatter curve is. A value of the Hill coefficient greater than 1 indicates positive cooperativity that is manifested as a steep concentration-response curve. The greater value of the Hill coefficient the steeper curve is. In contrast to the Black et al. slope factor, the proposed implementation of the Hill coefficient (Eq. 10) does not affect the centre or asymptotes of the hyperbola describing concentration-response curves. Therefore, the Hill coefficient does not affect relationships among these three parameters of the OMA, K_A_, E_MAX_ and τ. This makes OMA (Eq. 11) more practical in many ways.

### The general concept of the operational model of agonism

In general, OMA consists of two functions. One function describes the binding of an agonist to a receptor as the dependence of the concentration of agonist-receptor complexes [RA] on the concentration of an agonist [A]. The second function describes the dependence of functional response (E) to an agonist on [RA]. OMA expresses the dependence of E on [A].

### Rectangular hyperbolic OMA

#### Definition of OMA

In the simplest case, when both binding and response functions are described by rectangular hyperbola (Supplementary Information Figure S1), the resulting function is also a rectangular hyperbola. For example, in a bi-molecular reaction, the dependence of ligand binding to the receptor [RA] is described by Eq. 1 where [A] is the concentration of ligand and K_A_ is its equilibrium dissociation constant that represents the concentration of ligand at which half of the total number of receptors, R_T_, is bound to the ligand and the other half of the receptors is free.

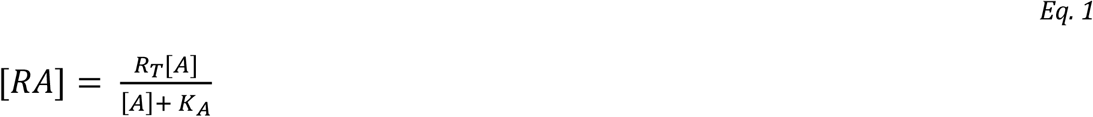

If the bound ligand is an agonist, it activates the receptor and produces functional response E. Response as a function of agonist binding (agonist-receptor complexes [RA]) is given by Eq. 2.

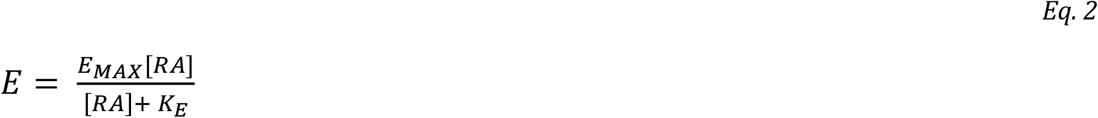

Where E_MAX_ is the maximum possible response of the system and K_E_ is the value of [RA] that elicits a half-maximal effect. Various agonists produce a functional response of different strengths. The OMA was postulated to introduce the “transducer ratio” τ that is given by Eq. 3.

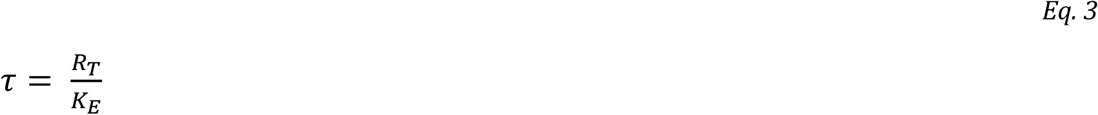

The substitution of Eq. 2 with Eq. 1 and Eq. 3 gives Eq. 4.

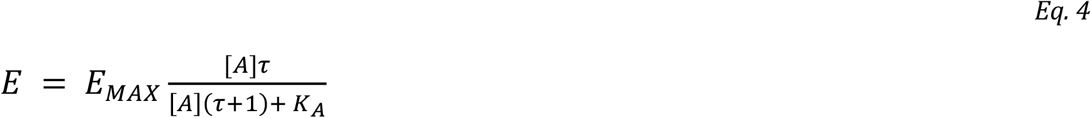

#### Analysis of OMA

Eq. 4 is the equation of OMA(Black and Leff, 1983). It has three parameters: The equilibrium dissociation constant of agonist (K_A_) at the response-inducing state of the receptor that is specific to a combination of ligand and receptor. The maximal possible response of the system (E_MAX_) is specific to the system. And the “transducer ratio” (τ) that is specific to a combination of ligand and system. Eq. 4 is a rectangular hyperbola with the horizontal asymptote, the observed maximal response to agonist A (E’_MAX_), given by Eq. 5.

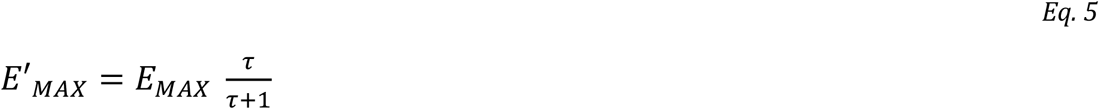

A more efficacious agonist (having a high value of parameter τ) elicits higher E’_MAX_ than less efficacious agonists (having a low value of parameter τ). Thus, τ is actually operational efficacy. The relationship between parameter τ and E’_MAX_ is hyperbolic meaning that two highly efficacious agonists (e.g., τ values 10 and 20) differ in E’_MAX_ values less than two weak agonists (e.g., τ values 0.1 and 0.2).

In Eq. 4, the concentration of agonist A for half-maximal response (EC_50_), is given by Eq. 6.

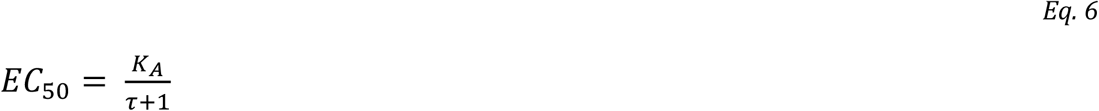

According to Eq. 6, for τ > 0, the EC_50_ value is always lower than the K_A_ value. The K_A_ to EC_50_ ratio is greater for efficacious agonists than for weak agonists. Similarly to E’_MAX_, the relationship between parameter τ and EC_50_ is hyperbolic. In contrast to E’_MAX_ values, the ratio K_A_ to EC_50_ ratio is more profound for two highly efficacious agonists (e.g., τ values 10 and 20) than for two weak agonists (e.g., τ values 0.1 and 0.2).

#### Limitations of OMA

The OMA has several weak points. The major drawback of OMA is the lack of physical basis of the agonist equilibrium dissociation constant K_A_. Eq. 2 assumes that agonist binding [AA] refers to agonist binding to the conformation from which the functional response is initiated. The agonist binding to the receptor in an inactive conformation is not observed in the response (K_E_→∞; τ = 0) In the radioligand binding experiments agonists bind to all receptor conformations including the inactive ones. For various reasons the receptors in the conformation activating functional response may be scarce or absent from radioligand binding experiments. Then it may be impossible to determine the KA value in the radioligand binding experiments. All three parameters of OMA (E_MAX_, K_A_ and τ) are interdependent (Jakubík et al., 2019). To fit Eq. 4 to the experimental data one of the parameters must be fixed. Thus, the maximal response of the system E_MAX_ has to be determined before fitting Eq. 4. It can be achieved by comparing the functional response to a given agonist in a system with a reduced population of receptors by irreversible alkylation(Furchgott, 1966). Another limitation of the OMA is that the shape of the functional response is a rectangular hyperbola.

### Non-hyperbolic OMA

#### Definition of non-hyperbolic OMA

In practice, concentration-response curves steeper or flatter than the ones described by Eq. 4 are observed. In such cases, Eq. 4 does not fit experimental data. As stated by the authors, Eq. 7 was devised for non-hyperbolic dependence of functional response on the concentration of agonist(Black et al., 1985).

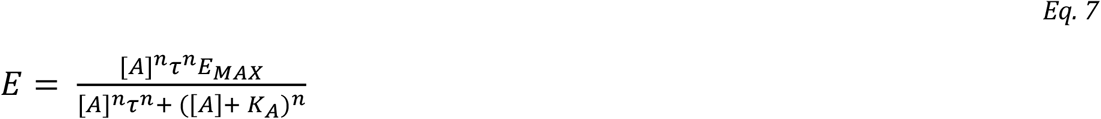

#### Analysis of non-hyperbolic OMA

Introduced power factor **n** changes the slope and shape of the functional-response curve (Supplementary information Figure S3). Nevertheless, Eq. 7 as a mathematical function has rectangular asymptotes: The horizontal asymptote ([A]→±∞) E’_MAX_ is given by Eq. 8 and the vertical asymptote (E→±∞) is equal to -K_A_/(τ+1) (Supplementary Information Eq. S2 and Eq. S3). From Eq. 7, the EC_50_ value is given by Eq. 9.

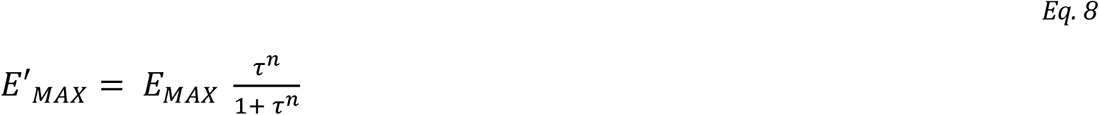

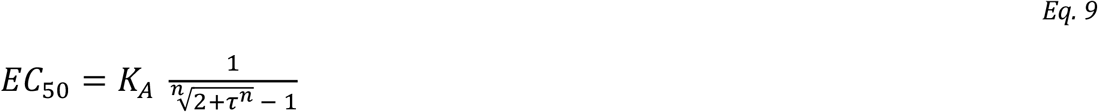

Evidently, the introduced slope factor **n** affects both the observed maximal response E’_MAX_ and the half-efficient concentration of agonist EC_50_ (Figure 1A and C). The influence of the slope factor on E’_MAX_ is **bidirectional** (Supplementary Information Table S1, Figure S2). For operational efficacies τ > 1, an increase in the value of the slope factor increases E’_MAX_. (Figure 1A and Figure 2A blue lines). For operational efficacies τ < 1, an increase in slope factor decreases E’_MAX_ (Figure 1C and Figure 2A yellow lines). The effect of slope factor on E’_MAX_ is the most eminent for low values of operational efficacy τ, making the estimation of model parameters of weak partial agonists impractical. Imagine full agonist τ=10 and very weak agonist τ=0.1. For n=1: Full agonist E’_MAX_ is 90 % and weak agonist E’_MAX_ is 10 % of system E_MAX_. For n=2: Full agonist E’_MAX_ is 99 % (one-tenth more) and weak agonist E’_MAX_ is just 1 % (ten times less).

**Figure 1.**
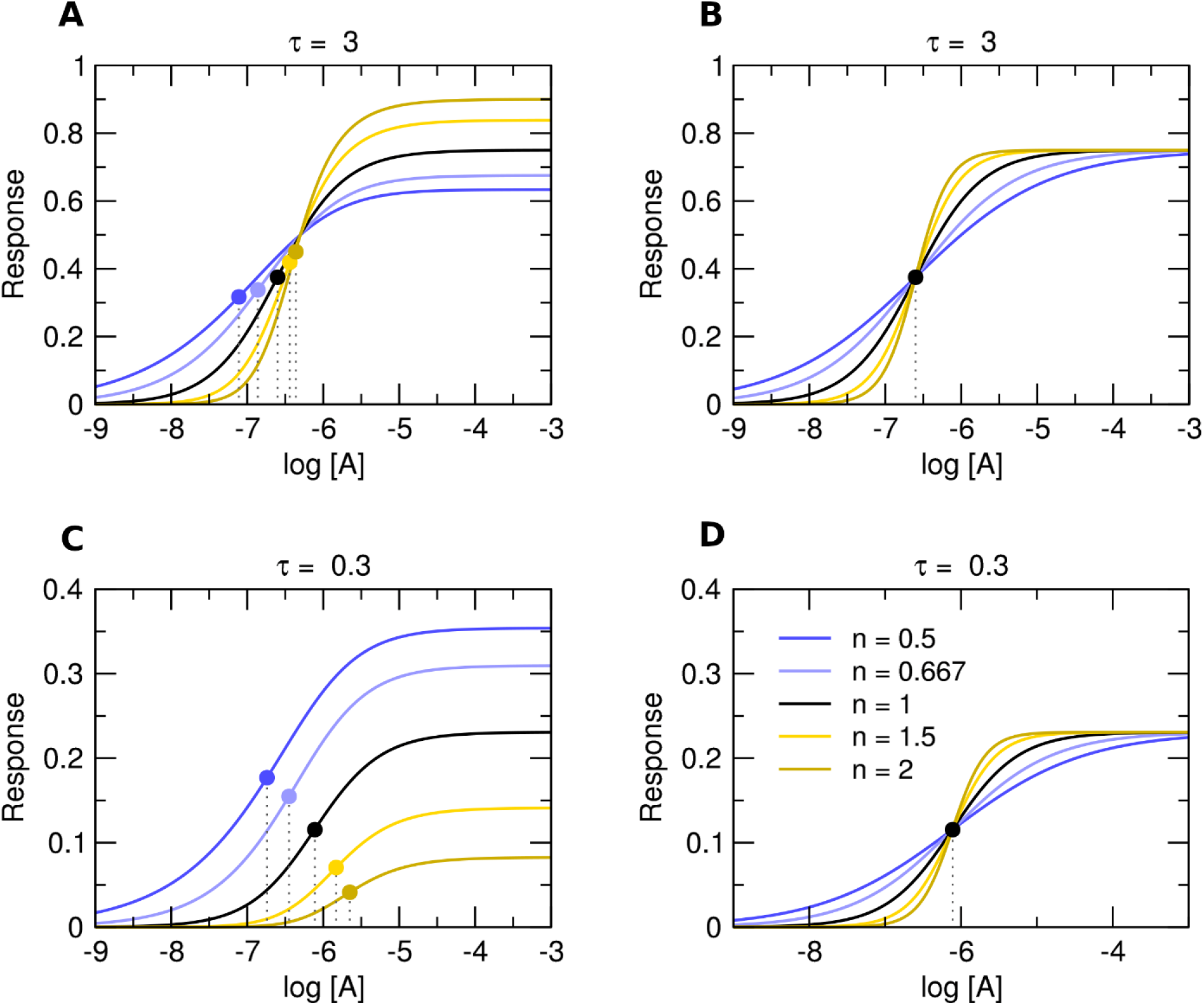
Theoretical concentration curves. Theoretical response curves according to Eq. 7, left and Eq. 11, right. Simulation parameters: E_MAX_ = 1; τ = 3 (top) or τ = 0.3 (bottom); K_A_ = 10^-6^ M. Values of slope factors are listed in the legend.

**Figure 2.**
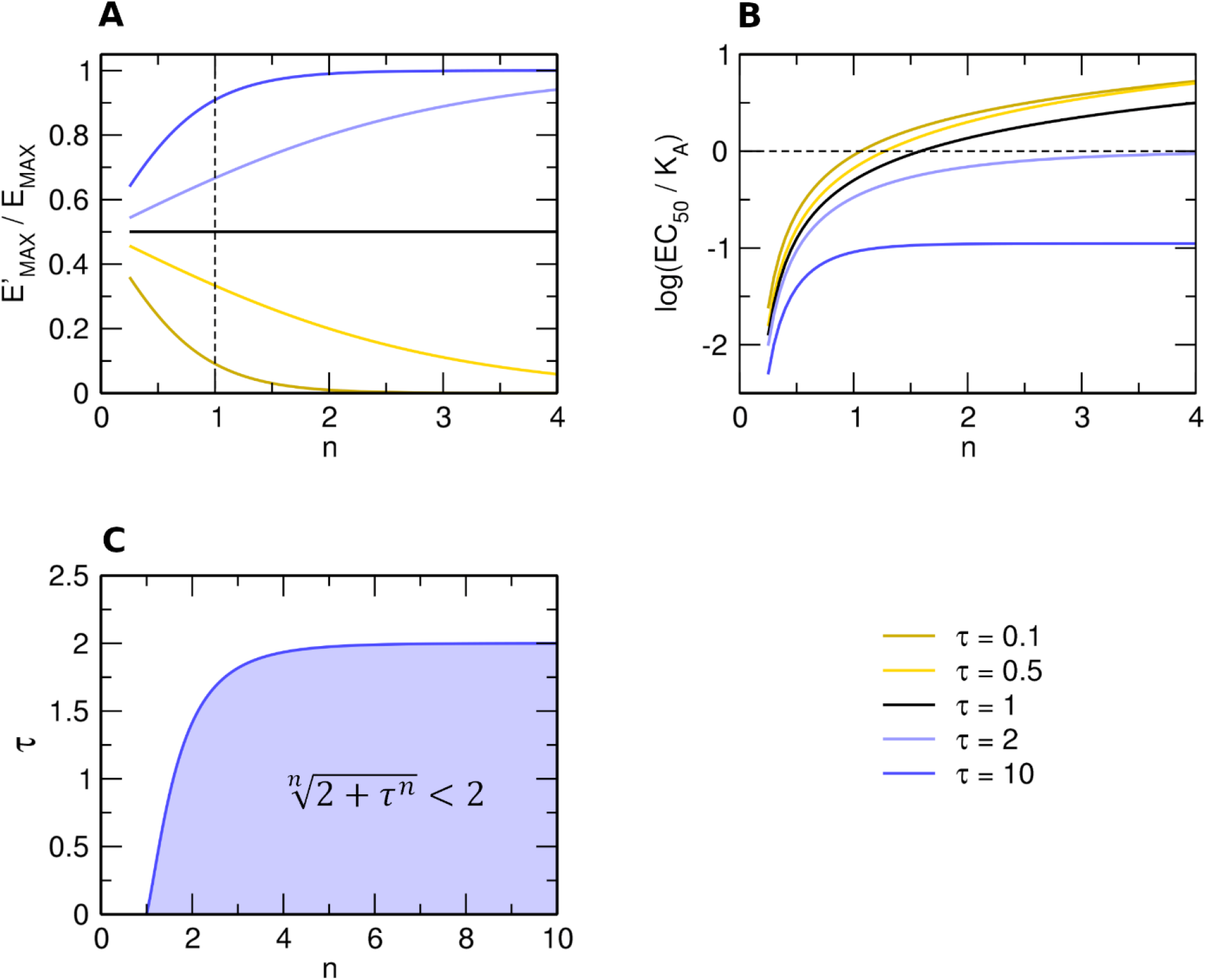
Analysis of Black & Leff equation (Eq. 7) A, Dependency of observed E’_MAX_ to system E_MAX_ ratio (ordinate) on slope factor n (abscissa) and operational efficacy τ (legend). B, Dependency of EC_50_ to K_A_ ratio (ordinate) on slope factor n (abscissa) and operational efficacy τ (legend). C, Inequality plot of slope factor **n** (abscissa) and operational efficacy τ (ordinate) yielding half-efficient concentration EC_50_ greater than equilibrium dissociation constant K_A_.

An increase in the value of the slope factor increases the EC_50_ value (Figure 2B). Again, the effect of the slope factor on the EC_50_ value is more eminent at low values of operational efficacy τ (red lines). Paradoxically, any combination of operational efficacy τ and slope factor fulfilling the inequality in Figure 2C (blue area) results in EC_50_ values **greater** than K_A_ (e.g., Figure 1C, yellow lines). For example, EC_50_ > K_A_ applies if τ = 0.5 and n > 1.6, or if τ = 1 and n > 1.6, or when τ = 1.5 and n > 2.15. The upper asymptote of inequality is 2. Thus, the possibility of EC_50_ > K_A_ applies to τ < 2.

The operational efficacy τ may be also considered as a measure of “receptor reserve”. In a system with a relatively small capacity of a functional response output, the full agonist reaches its maximal response before reaching full receptor occupancy. Thus, the agonist EC_50_ value is lower than its affinity for the receptor. The smaller the occupancy fraction needed for the full response to a given agonist the greater is difference between agonist EC_50_ and K_A_ values. According to OMA (Eq. 2), the relation between EC_50_ and K_A_ is described by Eq. 6. The greater value of operational efficacy τ, the smaller EC_50_ value and the greater the difference from K_A_. Thus, the value of operational efficacy τ is a measure of the receptor reserve of a given agonist in a given system. In a system with a large capacity of functional output, agonists do not have a receptor reserve and must reach full receptor occupancy to elicit a full signal. In such a system, even full agonists have small operational efficacies. Thus, the parameter τ is specific to a combination of ligand and system.

Nevertheless, for agonists that elicit at least some response in a given system, the parameter τ must be greater than 0. Then according to Eq.6 of the operational model of agonism, the EC_50_ value must be smaller than the K_A_ value. In principle, the EC_50_ value greater than the K_A_ can be achieved only by some parallel mechanism that increases the apparent K’_A_ provided that a ratio of K’_A_ to K_A_ is greater than EC_50_ to K_A_. For example, such a mechanism may be negative allosteric modulation of agonist binding or non-competitive inhibition of functional response.

#### Limitations of the non-hyperbolic OMA

Besides all limitations of the hyperbolic OMA, the non-hyperbolic version of OMA has additional drawbacks. The most important is the lack of mechanistic background for factor **n**. Exponentiation of agonist concentration [A] to power factor **n** results in non-hyperbolic functional-response curves. Importantly, as shown above, exponentiation of operational efficacy τ to power factor **n** breaks the logical relationship between observed maximal response E’_MAX_ and operational efficacy τ. That, as it will be shown later, impedes the correct estimation of τ and K_A_ values.

### OMA with Hill coefficient

#### Definition of OMA with Hill coefficient

The hill coefficient may serve as an alternative slope factor in the OMA. Hill equation incorporates the Hill coefficient as a slope factor to rectangular hyperbola (Gesztelyi et al., 2012). The major advantage of the Hill coefficient as a slope factor is that it allows for a change in the eccentricity (vertices) of the hyperbola-like curves without changing the centre (EC_50_) and asymptotes (E’_MAX_) (Supplementary Information Figure S3). The Hill–Langmuir equation published in 1910 can be formulated as Eq. 10.

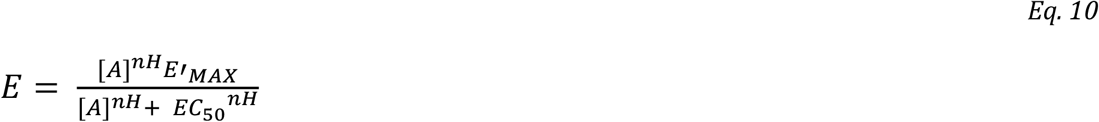

Where nH is the Hill coefficient. Substitution of E’_MAX_ by Eq. 5 and EC_50_ by Eq. 6 gives Eq. 11.

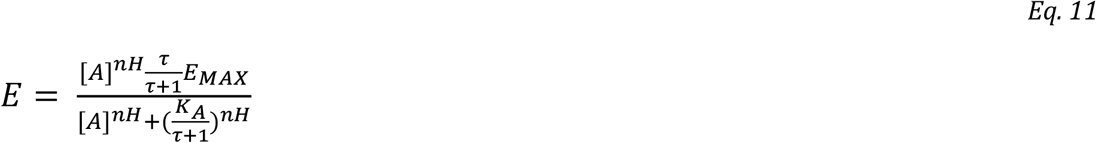

As expected, the Hill coefficient does not influence the maximal observed response E’_MAX_ or halfefficient concentration of agonist EC_50_ (Figure 1B and D). Eq. 11 was suggested as suitable for the analysis of functional responses displaying symmetrical response curves(Roche et al., 2016).

#### Implications of OMA with Hill coefficient

Analysis of the OMA with slope factor by Black et al. (Eq. 7) have shown that the slope factor **n** has a bidirectional effect on the relationship between the parameters E’_MAX_ and τ. and that the slope factor **n** affects the relationship between the parameters EC_50_ and K_A_. In contrast, in Eq. 11 neither value of E’_MAX_ nor the value of EC_50_ is affected by the Hill coefficient (Figure 1B and D). The parameters E’_MAX_ and EC’_50_ can be readily obtained by fitting Eq. 10 to the single concentration-response data.

#### Limitations of OMA with Hill coefficient

The major criticism of the Hill equation is its parsimonious character. It is relatively simple and its parameters are easy to estimate. However, as a model, it is just an approximation. In an experiment, the slope of the concentration-response curve different from unity may be a result of the parallel signalling mechanism providing feedback or allosteric cooperativity. E.g., positive cooperativity results in steep concentration-response curves and negative cooperativity results in flat concentration-response curves.

### OMA of allosteric systems

The simplest scenario leading to variation in the slope of concentration-response curves is allosteric interaction between two molecules of agonist, e.g., in a ligand-gated channel or dimeric GPCR(Jakubík and Randáková, 2022). Positive cooperativity among agonist molecules results in steep functional-response curves and negative cooperativity results in flat functional-response curves. As the mode of cooperativity is the property of a given agonist, the slope of the functional-response curve may vary among agonists. Slope factor **n** in Eq. 7 is deemed the property of the system and thus the same for all agonists. In the attempt to keep slope factor **n** constant in allosteric systems, the second slope factor **m** was introduced to Eq. 7 resulting in Eq. 12 (Gregory et al., 2020):

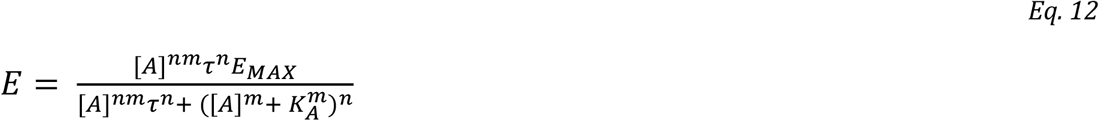

Where factor **n** links agonist concentration to the slope of the functional-response curve and factor **m** links agonist concentration to the slope of the receptor-binding curve. As the slope factors are interdependent one of them must be predetermined. It is only possible to predetermine the slope of the binding curve **m**. When fitting Eq. 12 to the data, functional-response slope factor **n** should become the same for all agonists. Observed maximal response E’_MAX_ is still given by Eq. 8 but half-efficient concentration EC_50_ is given by Eq. 13.

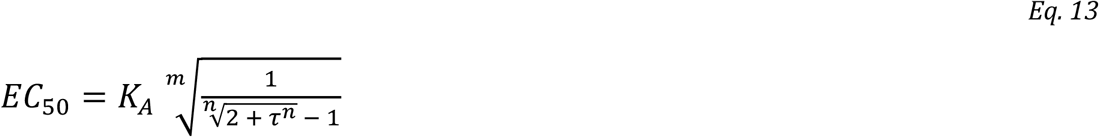

Similarly to Eq. 9, the effect of the binding slope factor **m** on the E_C50_ to K_A_ ratio depends on operational efficacy τ and functional response slope factor **n**. The effect of binding slope factor **m** depends on the radicand value. The m-th root is greater than the radicand when **m**>1 and radicand >1 or **m**<1 and radicant<1. The m-th root is smaller than the radicand when **m**<1 and radicand>1 or **m**>1 and radicand<1. Moreover, functional responses of allosteric systems may be not only steep or flat but also biphasic including bell-shaped(Jakubík and Randáková, 2022). Neither Eq. 7 nor Eq. 12 can accommodate such shapes and equations adequate to the mode of action are needed.

### OMA of substrate inhibition

Another common mechanism affecting the functional response to an agonist is substrate inhibition. In substrate inhibition, substrate concentration dependently inhibits the reaction. In the case of functional response to receptor activation, agonist-receptor complexes [RA] are the substrate (Figure 3). Functional response is given by Eq. 14.

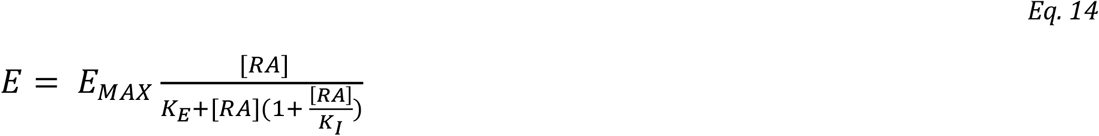

Where K_I_ is the inhibition constant, the concentration of [RA] that causes 50% inhibition of the functional response. After the substitution of Eq. 1 for [RA], Eq. 14 becomes Eq. 15 (Supplementary information Eq. S24).

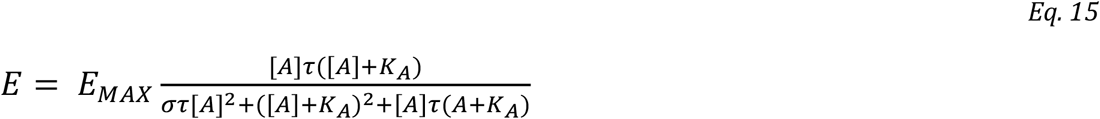

Where σ=R_T_/K_I_. The apparent maximal response E’_MAX_ is given by Eq. 16 (Supplementary Information Eq. S19).

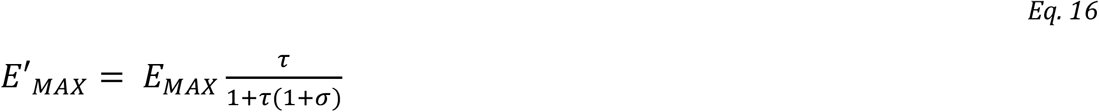

**Figure 3.**
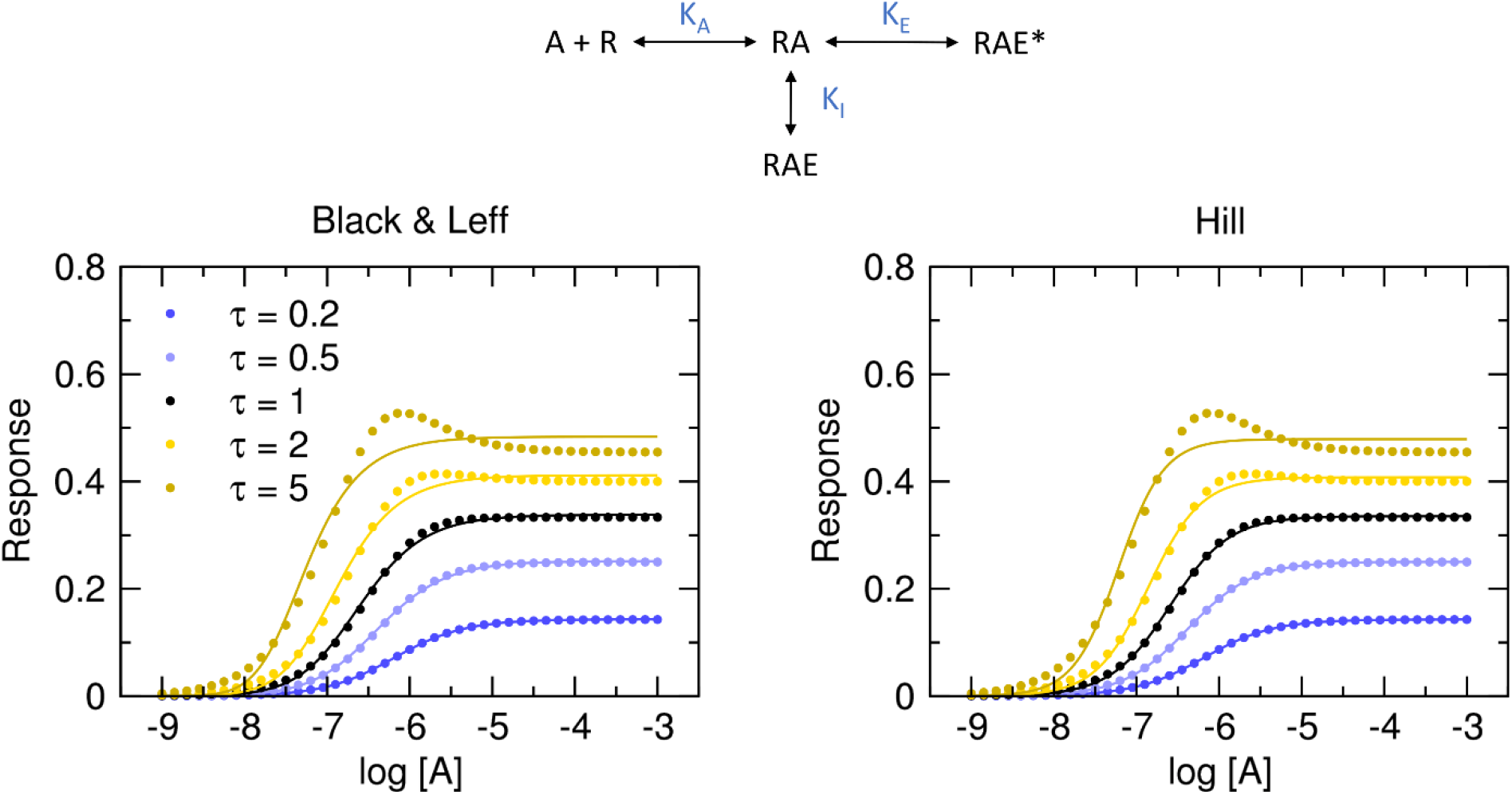
Substrate inhibition of functional response. Dots, functional response to an agonist (σ=1, E_MAX_=1, R_T_=1, K_A_ = 10^-6^ M) in the model of substrate inhibition according to Eq. 15. Values of operational efficacies τ are indicated in the legend. Full lines, left, Black & Leff equation (Eq. 7), right, Hill equation (Eq. 10) fitted to the data. Parameter estimates are in Table 1.

Thus, apparent operational efficacy τ’ is given by Eq. 17.

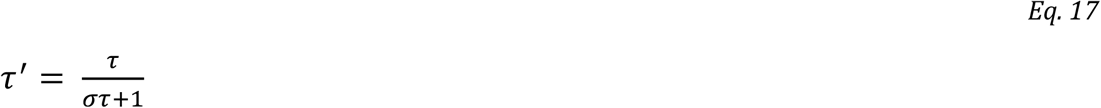

The EC_50_ value for the model of substrate inhibition is given by Eq. 18 (Supplementary information Eq. S27).

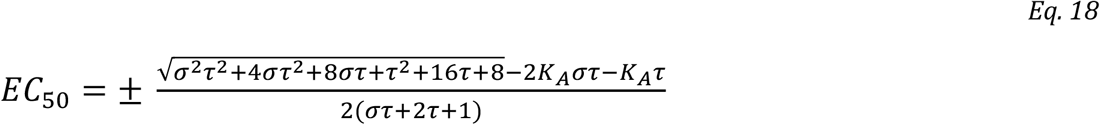

The Hill equation fits the data better than the Black & Leff equation. For strong inhibition (σ=10) curves of the functional response have a bell shape. Except for the bell-shaped curve, fitting of the Hill equation gives correct estimates of apparent maximal response E’_MAX_ and thus correct estimates of apparent operational efficacy τ’. In contrast, fitting the Black & Leff equation results in underestimated τ values and overestimated pK_A_ values. Stronger inhibition (greater value of σ) results in a larger overestimation of pKA.

Fitting Eq. 15 with fixed system EMAX to the model of functional response of substrate inhibition yields correct parameter estimates that are associated with a low level of uncertainty (Supplementary Information Figure S6).

### OMA of non-competitive inhibition

As shown in Figure 2, OMA with slope factor **n** allows for EC_50_ values higher than K_A_. The simplest mode of interaction that increases observed EC_50_ above K_A_ is non-competitive autoinhibition. Under non-competitive auto-inhibition, AR non-competitively blocks functional response by binding to effector E (Figure 4). It results in a concentration-dependent increase in EC_50_ and a decrease in E’_MAX_. Functional response is given by Eq. 19.

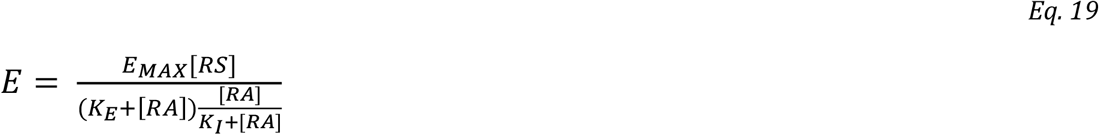

For K_I_ >0, after the substitution of Eq. 1 for [RA], Eq. 19 becomes Eq. 20 (Supplementary information Eq. S31).

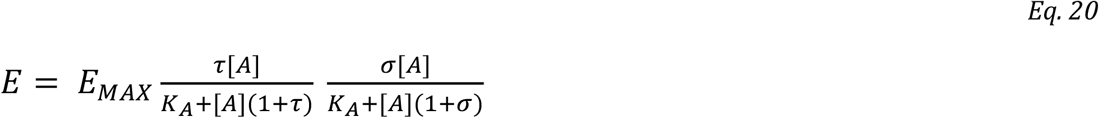

Where σ=R_T_/K_I_. The apparent maximal response E’_MAX_ is given by Eq. 21 (Supplementary Information Eq. S32).

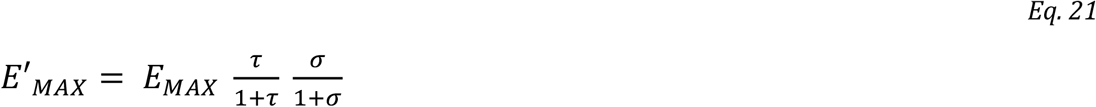

Thus, apparent operational efficacy τ’ is given by Eq. 22.

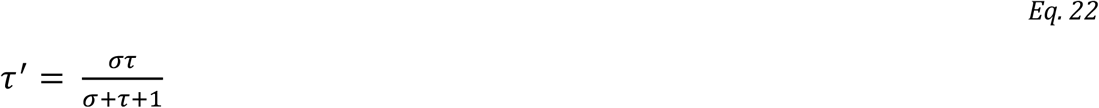

The EC_50_ value for the model of non-competitive auto-inhibition is given by Eq. 23 (Supplementary information Eq. S36).

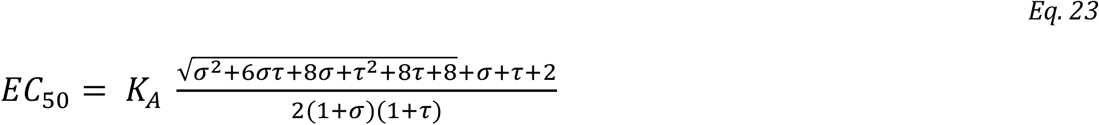

Non-competitive auto-inhibition decreases apparent maximal response E’_MAX_, increases observed half-efficient concentration EC_50_ and results in steep curves (Figure 4). The resulting concentration-response curve is asymmetric with a typical slope factor of about 1.2 regardless of values of K_I_ and K_E_. Both models fit well. Fitting Hill equation gives correct estimates of apparent maximal response E’_MAX_ and thus correct estimates of apparent operational efficacy τ’ (Table 2). In contrast, for K_I_≥1, fitting the Black & Leff equation results in underestimated values of τ and pKA.

**Figure 4.**
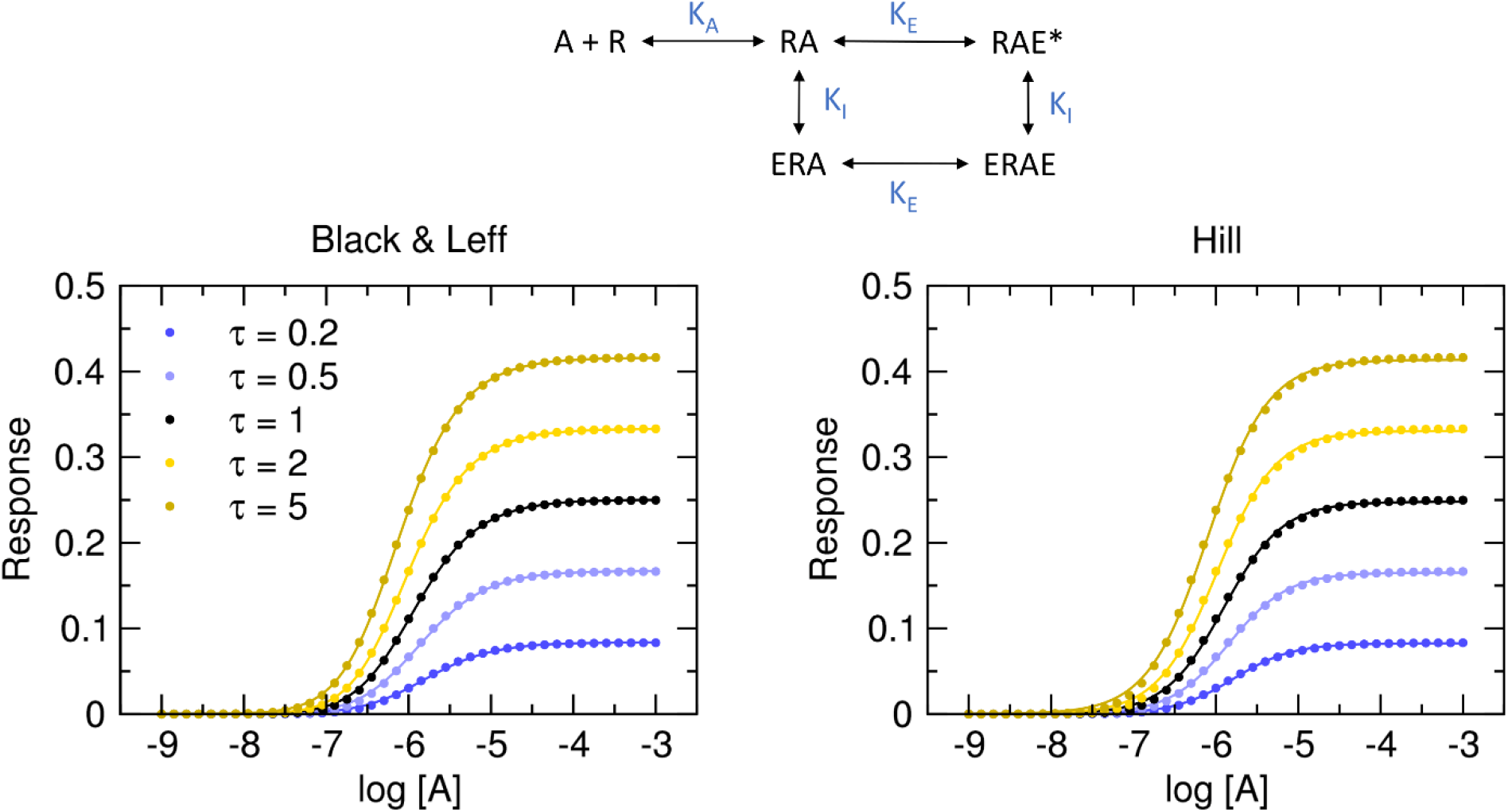
Non-competitive auto-inhibition of functional response. Dots, functional response to an agonist (K_I_=1, E_MAX_=1, R_T_=1, K_A_ = 10^-6^ M) in the model of noncompetitive autoinhibition according to Eq. 20. Values of operational efficacies τ are indicated in the legend. Full lines, left, Black & Leff equation (Eq. 7), right, Hill equation (Eq. 10) fitted to the data. Parameter estimates are in Table 2.

**Table 1.** Results of fitting Black & Leff and Hill equations to the model of substrate inhibition. Black & Leff (Eq. 7) and Hill (Eq. 10) were fitted to model data Eq. 15 with E_MAX_ fixed to 1.

| Model Eq. 15Eq. 17 |  |  |  | Black & Leff |  |  | Hill |  |  |  |
| --- | --- | --- | --- | --- | --- | --- | --- | --- | --- | --- |
| $\tau$ | $pK_A$ | $\sigma$ | $\tau'$ | $n$ | $\tau$ | $pK_A$ | $n_H$ | $EC_{50}$ | $E'_{MAX}$ | $\tau'$ |
| 0.2 | 6.00 | 1 | 0.167 | 1.09 | 0.194 | 6.17 | 1.04 | 6.18 | 0.142 | 0.166 |
| 0.5 | 6.00 | 1 | 0.333 | 1.23 | 0.411 | 6.38 | 1.11 | 6.38 | 0.250 | 0.333 |
| 1 | 6.00 | 1 | 0.500 | 1.45 | 0.628 | 6.65 | 1.21 | 6.60 | 0.335 | 0.504 |
| 2 | 6.00 | 1 | 0.667 | 1.84 | 0.824 | 7.02 | 1.36 | 6.85 | 0.408 | 0.689 |
| 5 | 6.00 | 1 | 0.833 | 2.74 | 0.976 | 7.57 | 1.59 | 7.22 | 0.474 | 0.901 |

**Table 2.** Results of fitting Black & Leff equations to the model of non-competitive auto-inhibition. Black & Leff (Eq. 7) and Hill (Eq. 10) equations were fitted to model data Eq. 20 with E_MAX_ fixed to 1.

| Model Eq. 20Eq. 22 |  |  |  | Black & Leff |  |  | Hill |  |  |  |
| --- | --- | --- | --- | --- | --- | --- | --- | --- | --- | --- |
| $\tau$ | pK<sub>A</sub> | K<sub>I</sub> | $\tau'$ | n | $\tau$ | pK<sub>A</sub> | n<sub>H</sub> | EC<sub>50</sub> | E'<sub>MAX</sub> | $\tau'$ |
| 0.2 | 6.00 | 1 | 0.091 | 1.33 | 0.166 | 5.91 | 1.21 | 5.78 | 0.083 | 0.090 |
| 0.5 | 6.00 | 1 | 0.200 | 1.46 | 0.333 | 5.98 | 1.22 | 5.84 | 0.165 | 0.199 |
| 1 | 6.00 | 1 | 0.333 | 1.58 | 0.499 | 6.06 | 1.22 | 5.91 | 0.248 | 0.330 |
| 2 | 6.00 | 1 | 0.500 | 1.71 | 0.666 | 6.15 | 1.21 | 5.99 | 0.331 | 0.494 |
| 5 | 6.00 | 1 | 0.714 | 1.55 | 0.804 | 6.15 | 1.18 | 6.10 | 0.414 | 0.706 |

Fitting Eq. 20 with fixed system E_MAX_ to the model of functional response of substrate inhibition yields correct parameter estimates that are associated with the low level of uncertainty only when correct initial estimates of τ and σ are given (Supplementary Information Figure S7 and S8). In the case of K_I_=5 (Supplementary Information Figure S9), estimates of operational efficacy τ and inhibition factor σ are swapped pointing to the symmetry of Eq. 20. This symmetry makes calculation of τ and σ impossible as any τ and σ combination resulting in an appropriate apparent efficacy τ’ (Eq. 22) corresponds well to a given functional-response data (Supplementary Information Figure S8). Fitting the Black & Leff equation (Eq. 7) yields wrong estimates of K_A_ and underestimated values of τ. Importantly, the extent of underestimation varies. The τ of 0.2 was underestimated by 17 %. The τ of 5 was underestimated 6-fold. In contrast, the calculation of apparent operational efficacies from the fitting of the Hill equation (Eq. 10) is very close.

### Signalling feedback

Signalling feedback occurs when outputs of a system are routed back as inputs (Figure 5). Negative feedback is a very common auto-regulatory (auto-inhibitory) mechanism in nature. Positive feedback also occurs in biology to propagate signals that would be otherwise dampened by other mechanisms. Functional response in the system employing feedback mechanisms is given by Eq. 24.

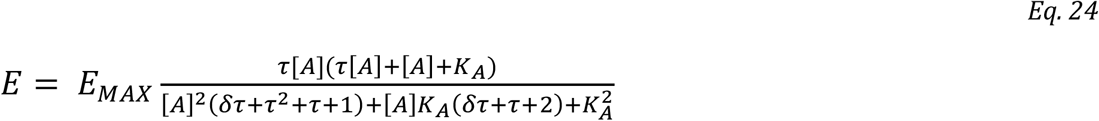

Where δ is a feedback factor. Values greater than 1 denote negative feedback. Values smaller than 1 denote positive feedback. The maximal response E’_MAX_ to an agonist with operational efficacy τ is related to the maximal response of the system E_MAX_ according to Eq. 25

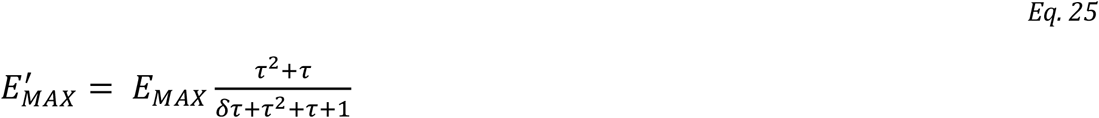

Negative feedback decreases the observed maximal response to an agonist E’_MAX_ while positive feedback increases it. And apparent operational efficacy τ’ of a given agonist can be calculated according to Eq. 26.

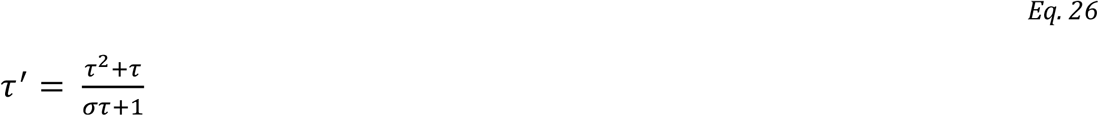

Furthermore, negative feedback decreases K_A_ to EC_50_ ratio while positive feedback increases it (Supplementary Information Eq. S54). For derivations see Supplementary Information Eq. S37 through Eq. S54. In general, negative feedback results in flat functional-response curves as with the increase in signal output proportionally more of an agonist is needed for the same increase of the signal. Conversely, positive feedback results in steep functional-response curves as signal output proportionally increases signal input (Supplementary Information Figure S9).

**Figure 5.**
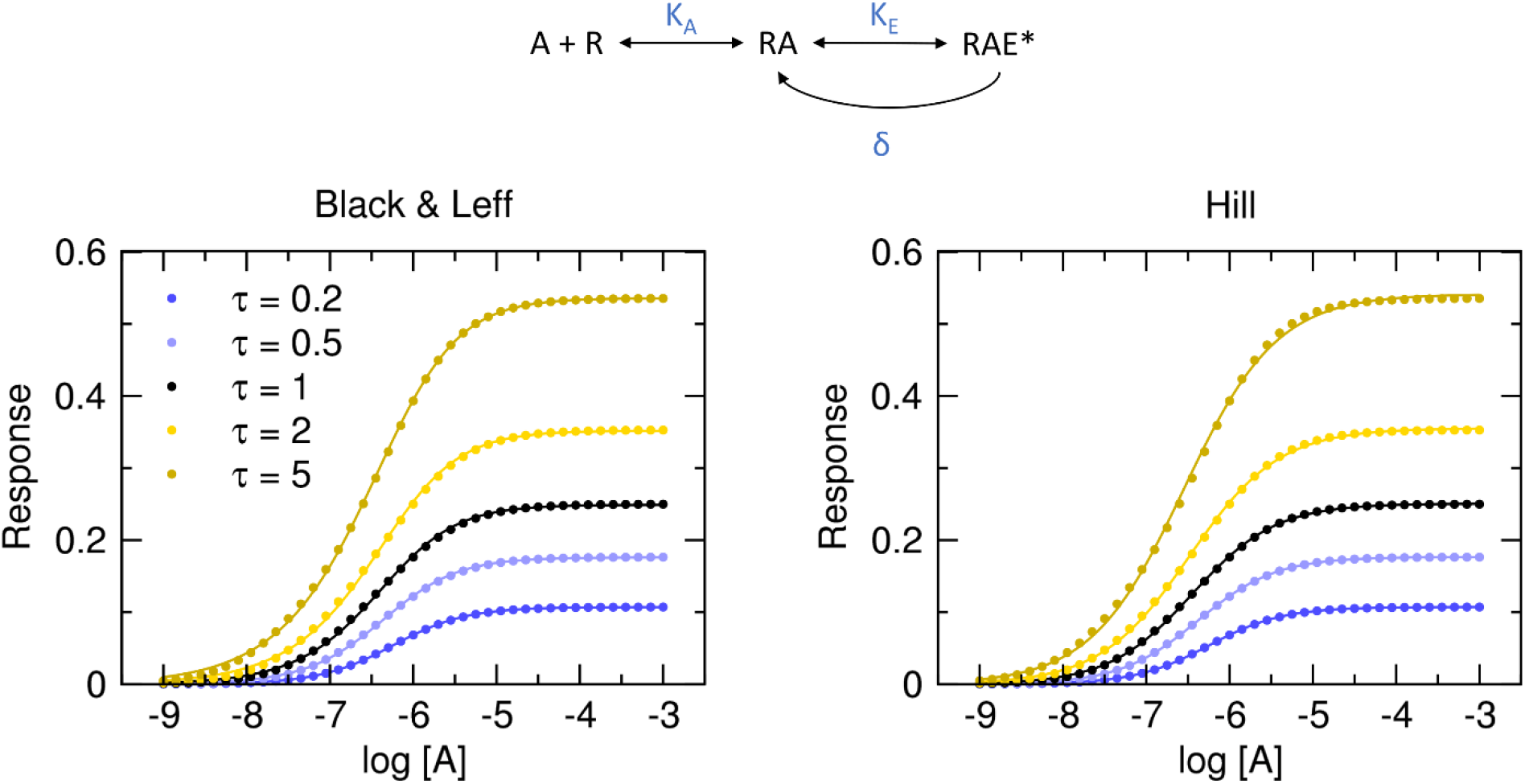
Functional response with signal feedback. Dots, functional response to an agonist (E_MAX_=1, R_T_=1, K_A_ = 10^-6^ M) in the model of the system with constant negative feedback (δ=5) according to Eq. 24. Values of operational efficacies τ are indicated in the legend. Full lines, left, Black & Leff equation (Eq. 7), right, Hill equation (Eq. 10) fitted to the data. Parameter estimates are in Table 3.

Importantly, in a system with constant negative feedback, like in Figure 5, the steepness of the curve depends on operational efficacy τ. Functional-response curves to agonists with high operational efficacy are flatter than the ones of agonists with low operational efficacy (Table 3).

**Table 3.** Results of fitting Black & Leff equations to the model of the system employing signalling feedback. Black & Leff (Eq. 7) and Hill (Eq. 10) equations were fitted to model data with E_MAX_ fixed to 1.

| Model Eq. 24 and Eq. 26 |  |  |  | Black & Leff |  |  | Hill |  |  |  |
| --- | --- | --- | --- | --- | --- | --- | --- | --- | --- | --- |
| $\tau$ | $pK_A$ | $\delta$ | $\tau'$ | $n$ | $\tau$ | $pK_A$ | $n_H$ | $pEC_{50}$ | $E'_{MAX}$ | $\tau'$ |
| 0.2 | 6.00 | 5 | 0.120 | 0.96 | 0.109 | 6.17 | 0.98 | 6.25 | 0.107 | 0.120 |
| 0.5 | 6.00 | 5 | 0.214 | 0.87 | 0.168 | 6.20 | 0.93 | 6.38 | 0.177 | 0.214 |
| 1 | 6.00 | 5 | 0.333 | 0.76 | 0.236 | 6.13 | 0.87 | 6.45 | 0.251 | 0.333 |
| 2 | 6.00 | 5 | 0.545 | 0.68 | 0.409 | 6.01 | 0.81 | 6.48 | 0.355 | 0.545 |
| 5 | 6.00 | 5 | 1.154 | 0.66 | 1.245 | 5.86 | 0.78 | 6.55 | 0.541 | 1.155 |

Fitting Eq. 24 with fixed system E_MAX_ to the model employing feedback mechanisms yields correct parameter estimates that are associated with the low level of uncertainty when correct initial estimates of τ and δ are given (Supplementary Information Figure S9 and S10). Fitting the Black & Leff equation (Eq. 7) yields wrong estimates of K_A_ and underestimated values of τ. Again, the extent of underestimation varies. The τ of 0.2 was underestimated by 67 %. The τ of 5 was underestimated 4-fold. In contrast, calculations of apparent operational efficacies from the fitting of the Hill equation (Eq. 10) are very close.

### Low receptor-expression systems

Eq. 2 and consequently Eq. 4 are valid only when [RA] is constant. That requires the number of RA complexes to be much greater than the number of available E. For example, this is not true in systems with low receptor-expression level. In a such system [RAE] as a function of [RA] is given by Eq. 27 (Supplementary Information Eq. S62).

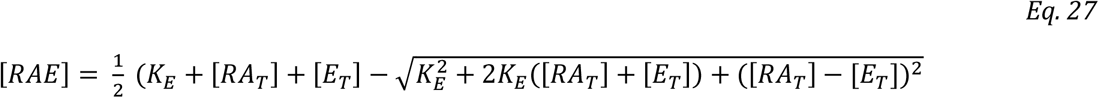

Where [RA_T_] denotes all receptor agonist complexes, free RA and RA in complex with E (RAE). [RAE] as a function of [A] (Supplementary Information Eq. S64) has only approximate solutions. Therefore, [RA_T_] values as a function of [A] were calculated according to Eq. 1 and used in Eq. 27 to model functional responses of the system with low receptor-expression level (Figure 6). The resulting curves are steep and asymmetric. With receptor pool depletion system reaches the maximum at lower agonist concentration which results in an increasing steepness and, in turn, in a curve asymmetry. The greater the operational efficacy is, the steeper and more asymmetric functional-response curves are (Table 4). However, the observed operational efficacy τ’ is equal to modelled operational efficacy.

**Figure 6.**
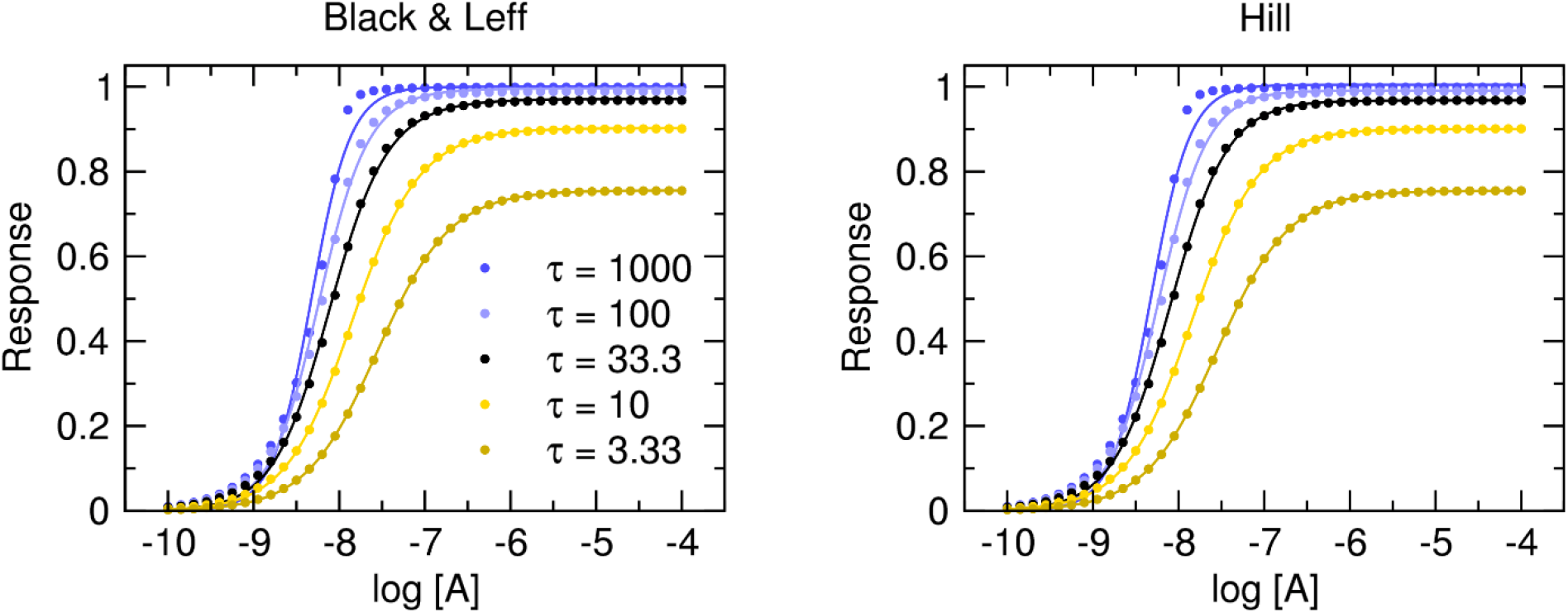
Functional response of the system with low receptor-expression level. Dots, functional-response data modelled in two steps. First, binding was calculated according to Eq. 1. Then resulting [RA] was used in Eq. 27. E_T_ = 10^-6^ M, R_T_ = 10^-5^ M, K_A_ = 10^-7^M. Values of operational efficacies τ are indicated in the legend. Full lines, left, Black & Leff equation (Eq. 7), right, Hill equation (Eq. 10) fitted to the data. Parameter estimates are in Table 4.

**Table 4.** Results of fitting Black & Leff equations to the model of the system with low receptor-expression level. Black & Leff (Eq. 7) and Hill (Eq. 10) equations were fitted to model data with E_MAX_ fixed to 1.

| Model Eq. 1 and Eq. 27 |  | Black & Leff |  |  | Hill |  |  |  |
| --- | --- | --- | --- | --- | --- | --- | --- | --- |
| $\tau$ | $pK_A$ | $n$ | $\tau$ | $pK_A$ | $n_H$ | $pEC_{50}$ | $E'_{MAX}$ | $\tau'$ |
| 1000 | 7.00 | 1.91 | 6701 | 4.50 | 1.89 | 8.32 | 1.00 | Inf. |
| 100 | 7.00 | 1.54 | 36.2 | 6.68 | 1.52 | 8.23 | 0.99 | 108 |
| 33.3 | 7.00 | 1.35 | 13.6 | 6.98 | 1.29 | 8.10 | 0.97 | 30.1 |
| 10 | 7.00 | 1.15 | 6.94 | 6.98 | 1.11 | 7.84 | 0.90 | 9.04 |
| 3.33 | 7.00 | 1.05 | 2.93 | 6.98 | 1.03 | 7.55 | 0.75 | 3.08 |

Fitting the Black & Leff equation (Eq. 7) to the model system with low receptor-expression level yields wrong estimates of K_A_ for extremely high operational efficacy (τ = 1000). Values of τ are far off for all modelled efficacies. While τ of 100 is overestimated more than 6-fold, the other τ values are underestimated up to 3-fold. Except for the τ value of 1000, operational efficacies calculated according to the Hill equation (Eq. 10) are less than 10 % off the model values.

## Discussion

The operational model of agonism (OMA)(Black and Leff, 1983) is widely used in the evaluation of agonism. The OMA characterizes a functional response to an agonist by the equilibrium dissociation constant of the agonist (K_A_), the maximal possible response of the system (E_MAX_) and the operational efficacy of the agonist (τ) (Eq. 4). To fit non-hyperbolic functional responses slope factor **n** was introduced to the OMA (Eq. 7) (Black et al., 1985). Analysis of the Black & Leff equation (Eq. 7) has shown that the slope factor **n** has a bidirectional effect on the relationship between the parameters E’_MAX_ and τ (Figure 1A versus C) and also affects the relationship between the parameters EC_50_ and K_A_.

Fitting Black & Leff equation (Eq. 7) to the models of substrate inhibition (Figure 3, Table 1) and non-competitive auto-inhibition (Figure 4, Table 2), signalling feedback (Figure 5, Table 3) and system with ow receptor-expression level (Figure 6, Table 4) resulted in wrong estimates of τ and K_A_ values. In the presented examples, the degree of over- or under-estimation of τ is not the same for all its values but depends on the value of τ, distorting relations among estimates of τ vales. In contrast, fitting the Hill equation (Eq. 10) to the model data gave more or less accurate estimates of apparent operational efficacies τ’ from which operational efficacies can be calculated (according to Eq. 17, Eq. 22, and Eq. 26), provided that mechanism of action is known.

Biased agonists stabilize specific conformations of the receptor leading to non-uniform modulation of individual signalling pathways (Lefkowitz, 2013). To measure an agonist bias, the parameters τ and K_A_ must be determined and log(τ/KA) values of tested and reference agonists compared at two signalling pathways (Kenakin et al., 2012). It is evident from the analysis of the Black & Leff equation, that as far as the EC_50_ value is dependent on parameters **n** and τ (Figure 1A and C), log(τ/K_A_) values cannot be compared to judge possible signalling bias unless the parameter **n** is equal to 1 for both tested and reference agonist.

Despite the dire effects of slope factor **n**, the Black & Leff equation is widely accepted (Christopoulos and El-Fakahany, 1999; Kenakin, 2012, 2017; Kenakin and Christopoulos, 2013; Keov et al., 2014; Luttrell et al., 2015; Stott et al., 2016; Burgueño et al., 2017). It even entered textbooks (Kenakin, 2014). Very little concern on factor **n** has risen. For example, Kenakin et al.(Kenakin et al., 2012) analysed in detail the effects of slope factor **n** on EC_50_ and τ to K_A_ ratio but did not deal with the bi-directional effect of **n** on τ nor proposed an alternative approach to avoid potential pitfalls. To force a proper shape on functional-response curves, Gregory et al.(Gregory et al., 2020) introduced the second slope factor into their OMA and operational model of allosterically modulated agonism (OMAMA) analysis making equations even more complex. So far, the greatest criticism of OMA was voiced by Roche et al. (Roche et al., 2016), noting that to accommodate the shape of theoretical curves Black & Leff equation tends to overestimate equilibrium dissociation constant K_A_ and operational efficacy τ and thus be misleading. They advocated for different expressions of operational models including OMA modified by the Hill coefficient in the case of symmetric concentration-response curves.

The fitting of the Hill equation (Eq. 10) to the functional response is straightforward and easier than fitting the Black & Leff equation. As shown in Figure 3 to Figure 6, the Hill equation fits well with various functional-response curves, often better than the Black & Leff equation. In contrast to the Black & Leff equation, the Hill equation gives correct estimates of maximal response to agonist E’_MAX_ and its half-efficient concentration EC_50_ as documented in Table 1 to Table 4. In the case of the Hill equation, neither value of E’_MAX_ nor the value of EC_50_ is affected by the Hill coefficient (Figure 1B and D). Therefore, biased signalling may be inferred from the comparison of the ratio of intrinsic activity (E’_MAX_/EC_50_) of tested agonist to the intrinsic activity of reference agonist at two signalling pathways as in the case of the Hill equation the E’_MAX_/EC_50_ ratio is equivalent to τ / K_A_ ratio (Ehlert et al., 1999; Griffin et al., 2007).

Further, if needed, apparent operational efficacy τ’ can be calculated from known E’_MAX_ values and known maximal response of the system E_MAX_. From the relationship between τ’ and EC_50_ (Jakubík et al., 2019, 2020) the mechanism of functional response can be inferred by comparison to explicit models. Four examples of explicit models are given above. Subsequently, equilibrium dissociation constant K_A_ and operational efficacy τ alongside the rest of the model-specific parameters can be obtained by fitting the explicit model to experimental data.

## Conclusions

Analysis of the Black & Leff equation has shown that i) The slope factor **n** has a bidirectional effect on the relationship between the parameters E’_MAX_ and τ. ii) The slope factor **n** affects the relationship between the parameters EC_50_ and K_A_. Fitting the Black & Leff equation gives wrong estimates of τ and K_A_ values when slope factor **n** differs from unity, limiting the use of the Black & Leff equation in the evaluation of signalling bias. Analysis of the Hill equation has shown that the Hill coefficient does not affect the relationship between the parameters E’_MAX_ and τ nor between the parameters EC_50_ and K_A_. Fitting the Hill equation to the concentration-response data gives good estimates of EC_50_ and E’_MAX_ values, which are suitable for further analysis of and signalling bias, e.g. using relative intrinsic activities.

## Acknowledgements

This research was funded by the Czech Academy of Sciences institutional support [RVO:67985823], the Grant Agency of the Czech Republic grant [23-04670S] and by the project National Institute for Neurological Research (Programme EXCELES, ID Project No. LX22NPO5107) - Funded by the European Union - Next Generation EU.

## Author contributions

JJ derived equations for theoretical models and wrote python code for model data analysis.

## Conflict of interest

The author declares no conflict of interest.

## Availability of data

The python code used in this study is available from the author upon request.

## Supplementary Information

### Hyperbola

A hyperbola is the set of points in a plane whose distances from two fixed points, called foci, has an absolute difference that is equal to a positive constant. It consists of two separate curves, called branches. Hyperbola is completely determined by its centre, vertices, and asymptotes. Points on the separate branches of the graph where the distance is at a minimum are called vertices. The midpoint between vertices is its centre. Hyperbola is asymptotic to certain lines drawn through the centre. Hyperbola which asymptotes form right angle is called rectangular or equilateral. The simplest rectangular hyperbola with coordinate axes as its asymptotes is reciprocal function y = 1/x (Figure S1, black). The reciprocal function can be geometrically translated by coefficients a, b, c, and d (Figure S1, green a=1, b=1, c=1, d=-1).

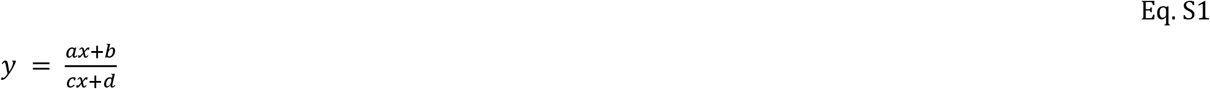

Where vertical and horizontal asymptotes are given by Eq. A2 and A3, respectively.

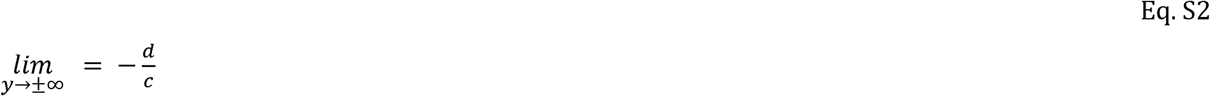

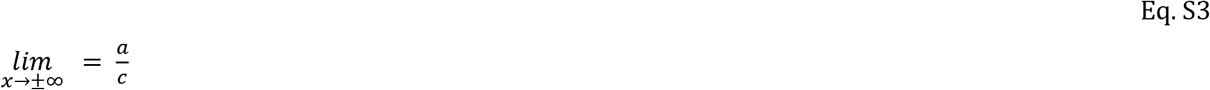

**Figure S1.**
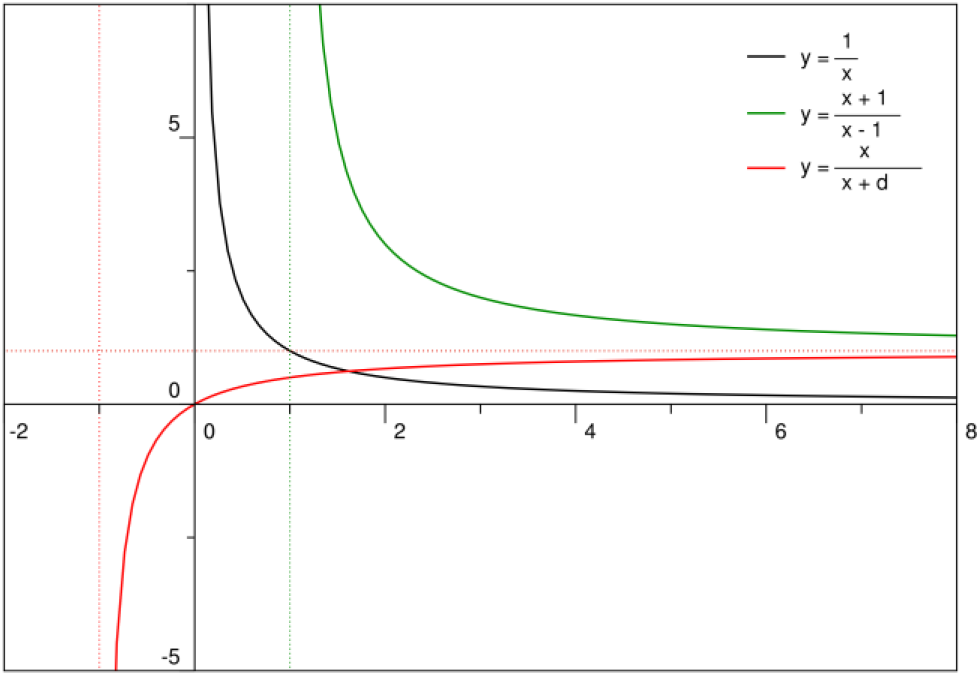
Examples of rectangular hyperbolas. Green, hyperbola with parameters a=1, b=1, c=1 and d=-1. Red, hyperbola with parameters a=1, b=0, c=1, d=1. Dashed lines, vertical and horizontal asymptotes of respective hyperbolas. Only right branches are shown.

A hyperbola with parameters a > 0, b = 0, c = 1 and d > 0 (e.g., Figure S1 red) is mass action equilibrium function.

### Power function

Exponentiation is a mathematical operation, written as b^n^, involving two numbers, the base b and the exponent n. b ≥ 0 can be risen to any value of n. b < 0 can be risen only to integers or fractions with odd denominator. Table S1 summarizes relationships between b^n^ and b values ≥ 0. Reciprocal relationships apply for b values < 0 as the power function is symmetric at the origin of axes. Importantly relationship between values of b^n^ and n are opposite in the range 0 < b < 1 to the range b > 1, resulting in the S-shape of the function (Figure S2).

**Table S1.** Behaviour of power function, relationship between b^n^ and b values.

| | $n = 0$ | $0 < n < 1$ | $n = 1$ | $n > 1$ |
| --- | --- | --- | --- | --- |
| $b = 0$ | $b^n = 1$ | $b^n = b$ | $b^n = b$ | $b^n = b$ |
| $0 < b < 1$ | $b^n = 1$ | $b^n > b$ | $b^n = b$ | $b^n < b$ |
| $b = 1$ | $b^n = 1$ | $b^n = b$ | $b^n = b$ | $b^n = b$ |
| $b > 1$ | $b^n = 1$ | $b^n < b$ | $b^n = b$ | $b^n > b$ |

**Figure S2.**
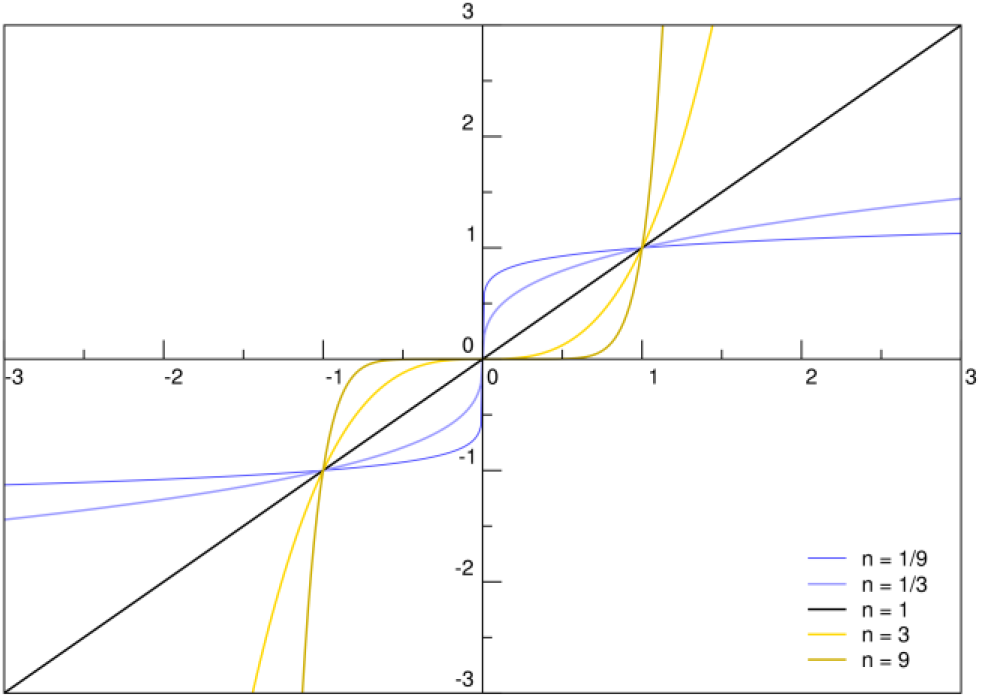
Examples of power functions. Curves of y = x^n^ for various values of exponent n. Each curve passes through the point (0, 0) because 0 raised to any power is 0 and through the point (1, 1) because number 1 raised to any power is 1. For n = 1, y = x because any number raised to the power of 1 is the number itself.

### Hyperbola and power function

Exponentiation of x in the hyperbola (Eq. A1) results in non-hyperbolic function (Eq. A4, Figure S3).

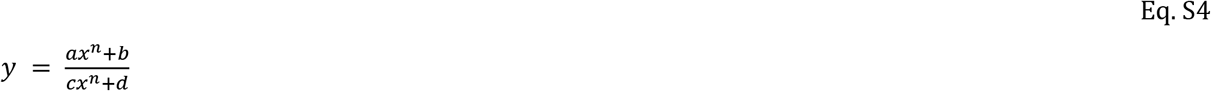

However, resulting function still possesses asymptotes given by Eq. A2 and A3 (Figure S3).

**Figure S3.**
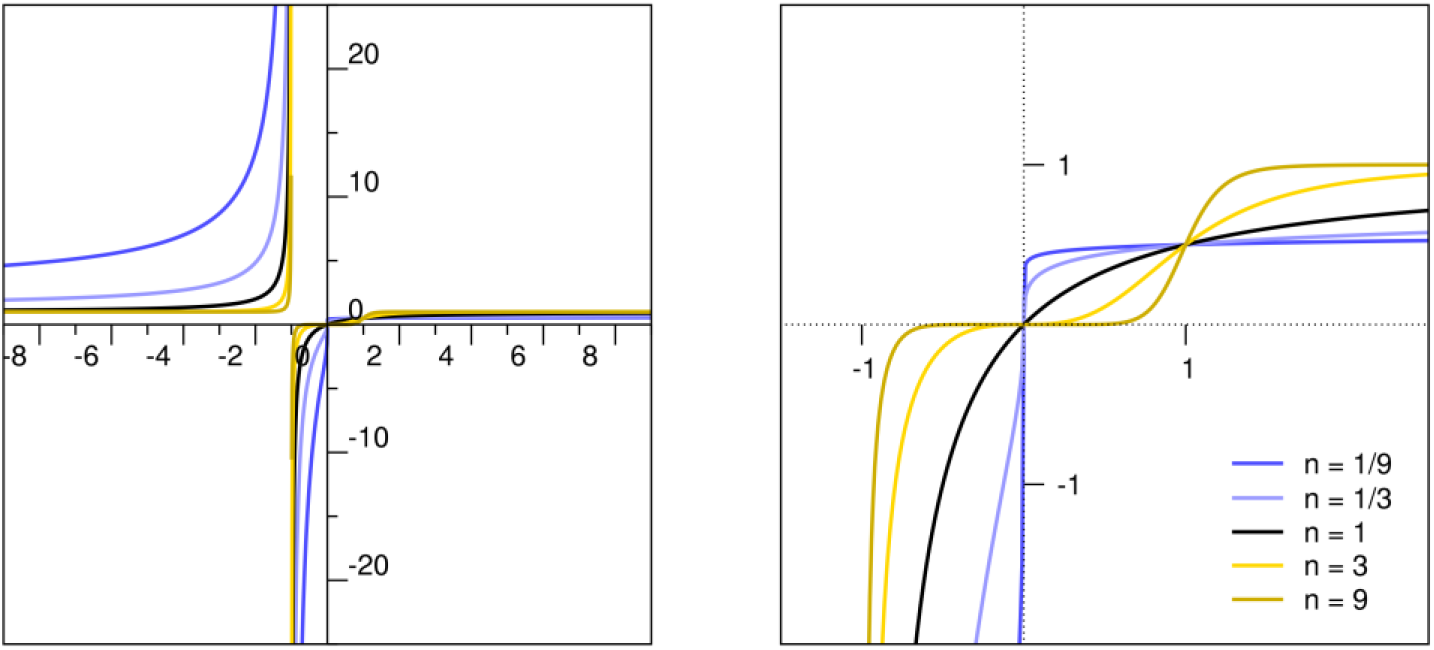
Examples of functions according to Eq. A4. Curves according to Eq. A4 for various values of exponent n. Left, both branches are shown. Right, detail of right branches. Black, for n=1 curve is rectangular hyperbola. Shades of blue, n < 1, and shades of yellow, n > 1, curves are S-shaped.

### OMA from scratch

Binding:

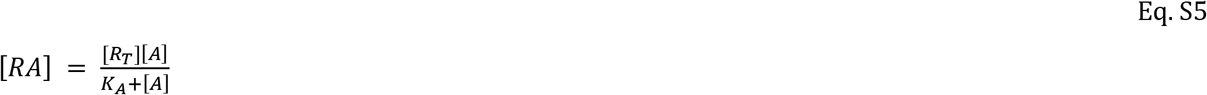

Functional response:

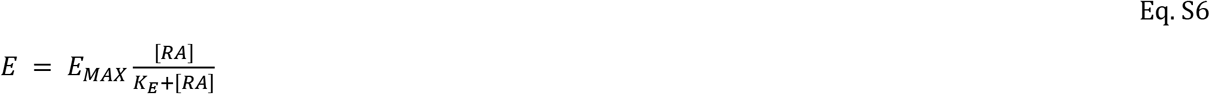

Substitution of [RA] by Eq. S5:

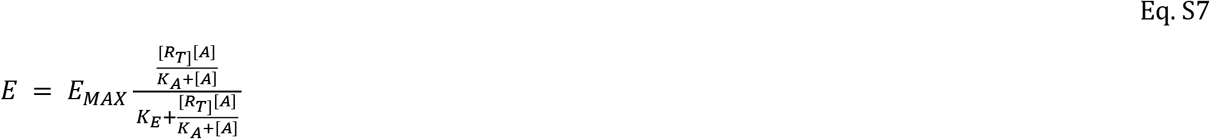

After rearrangement:

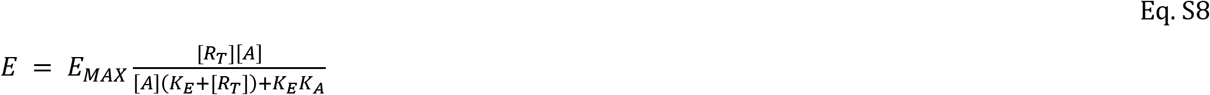

Division by K_E_ and substitution τ=[R_T_]/K_E_:

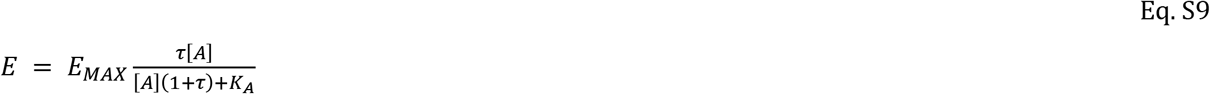

Apparent E’_MAX_ for [A]>>K_A_:

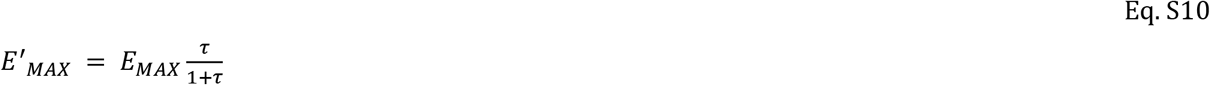

For half-efficient concentration EC_50_ Eq. S9 equals half of Eq. S10:

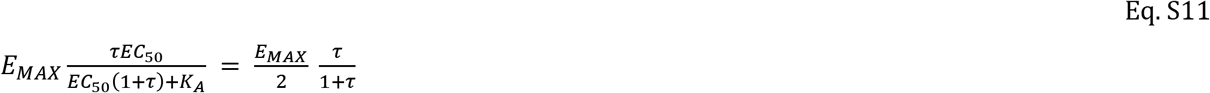

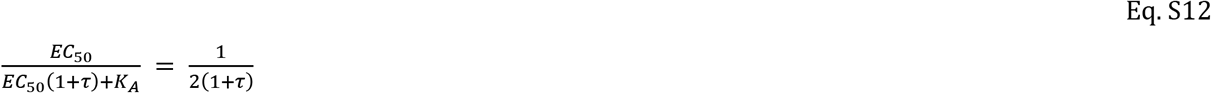

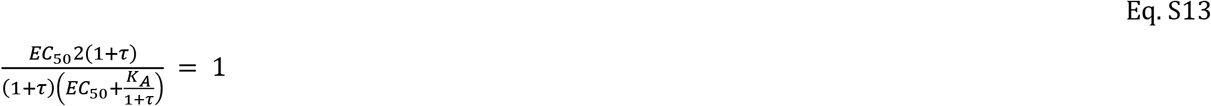

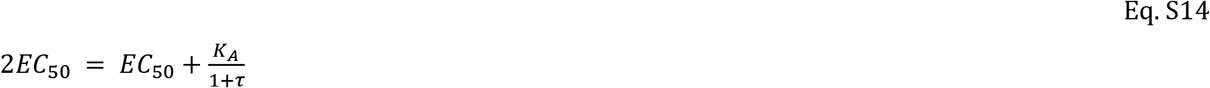

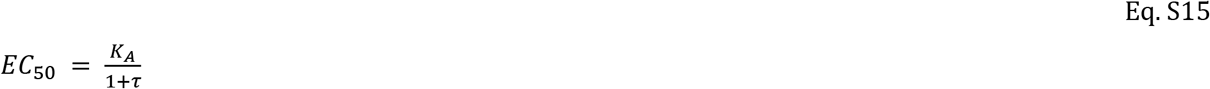

### Effect of changes in receptor concentration on functional response

**Figure S4.**
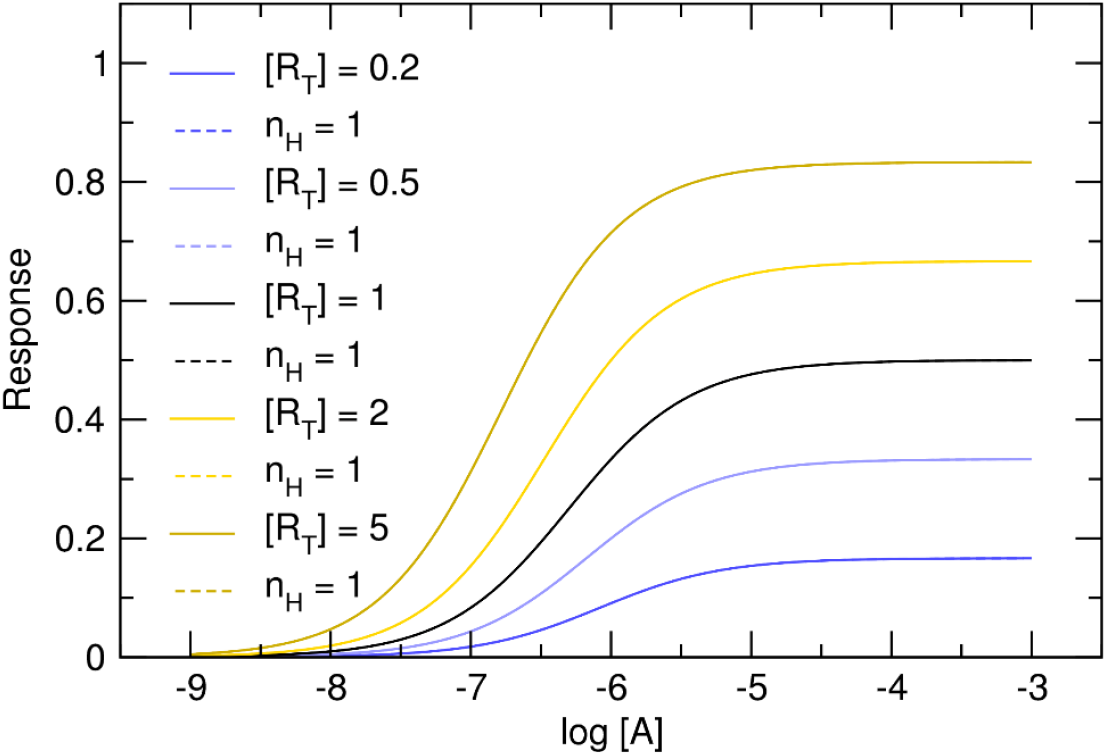
Effect of changes in receptor number on functional response. Curves according to OMA for various receptor concentrations ([R_T_]). E_MAX_ = 1, K_A_ = 10^-6^M, τ = 1, [R_T_] is in indicated in the legend.

### Effect of signal amplification on functional response

**Figure S5.**
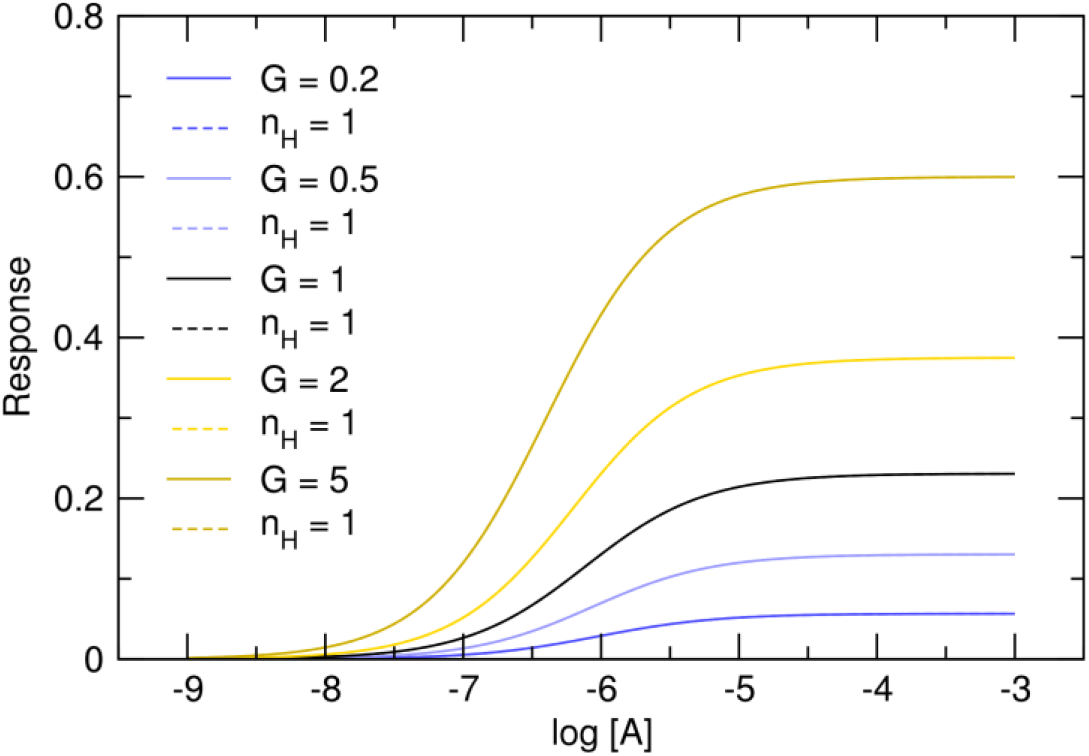
Effect of signal amplification on functional response. Curves according to OMA for various extent of signal amplification, gain factor (G) is indicated in the legend. E_MAX_ = 1, K_A_ = 10^-6^M, τ = 0.3, [R_T_] is in indicated in the legend.

### Substrate inhibition of functional response

In the case of substrate inhibition, functional response is given by Eq. S16.

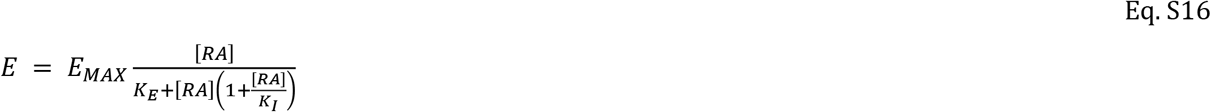

Substitution of Eq. S16 [RA] by binding function Eq. S5:

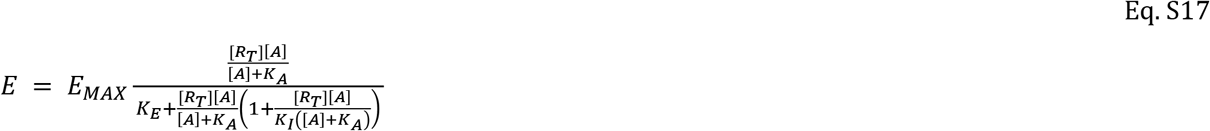

Apparent E’_MAX_ at [A]>>K_A_ is given by:

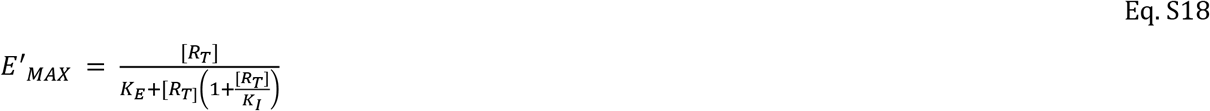

After division of Eq. S18 by K_E_ and substitution τ=[R_T_]/K_E_ and σ=[R_T_]/K_I_:

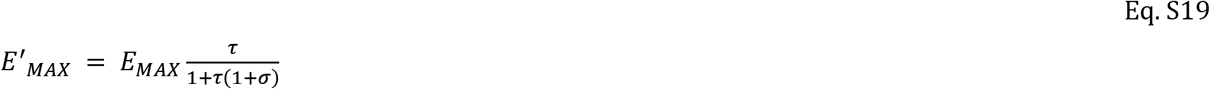

The apparent value of operational efficacy τ’ is given by equation Eq. S20.

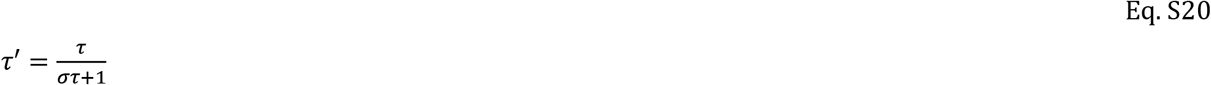

Denominator of Eq. S17 can be rearranged to:

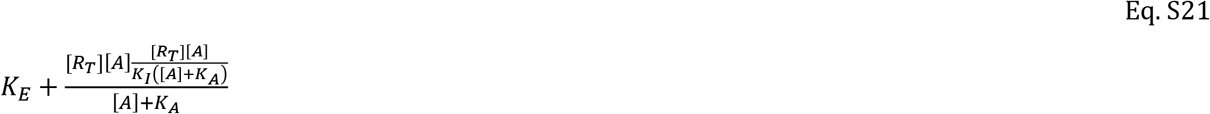

The Eq. S17 can be rearranged to Eq. S22.

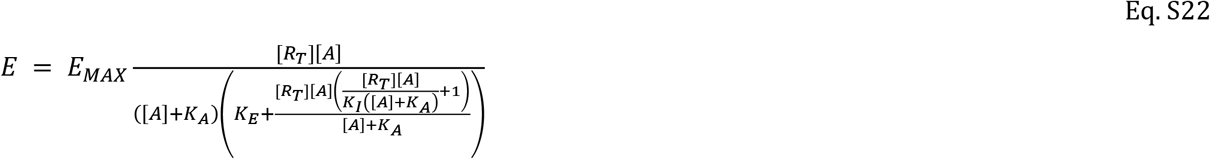

Eq. S22 after bracket multiplication and rearrangement gives Eq. S23.

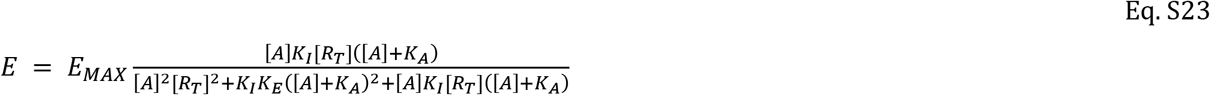

After division by K_E_ and K_I_ and substitution τ=[R_T_]/K_E_ and σ=[R_T_]/K_I_:

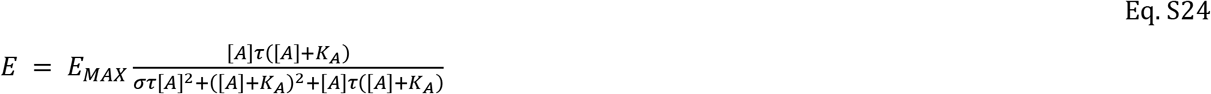

To determine half-efficient concentration EC_50_:

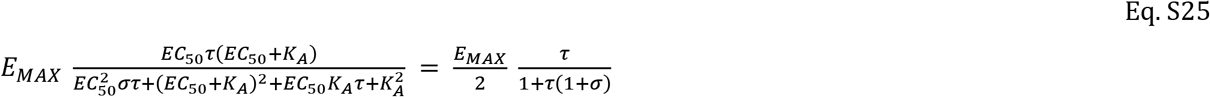

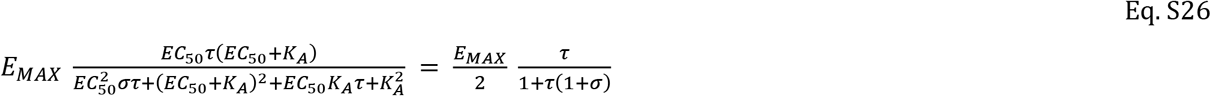

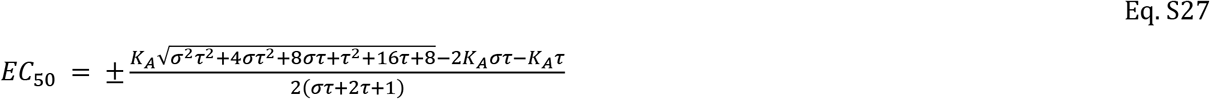

**Figure S6.**
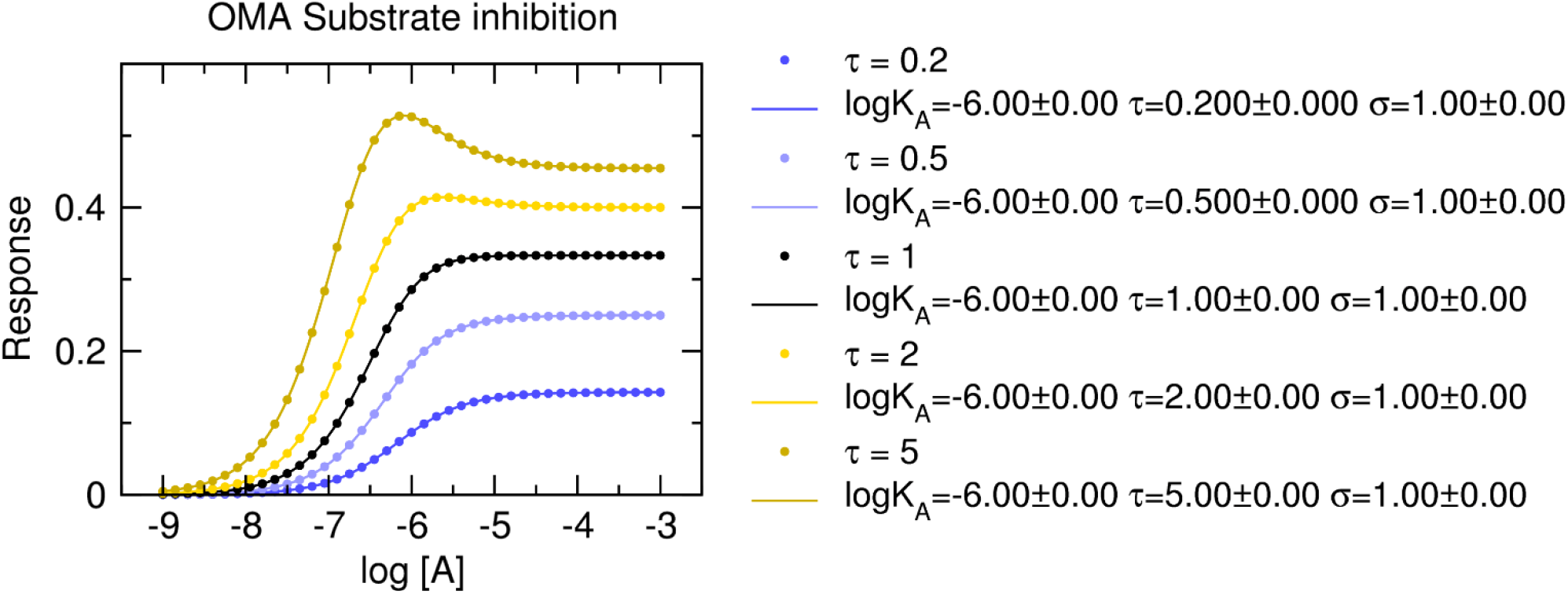
Fitting Eq. S24 to the model of substrate inhibition. Dots, functional-response data modelled according to Eq. S17. E_MAX_ = 1, K_A_ = 10^-6^M, σ = 1, [R_T_] =1. Values of operational efficvacy τ are indicated in the legend. Lines, fits of Eq. S24 to the model data. E_MAX_ was fixed to 1. Parameter estimates of the fits are indicated in the legend.

Parameter estimates are correct, associated with low level of uncertainty.

### Non-competitive auto-inhibition

In the case of non-competitive auto-inhibition, functional response is given by Eq. S28..

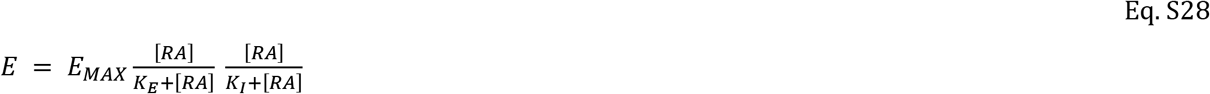

Substitution of the [RA] complexes in Eq. S28 with binding function Eq. S5 yields Eq. S29.

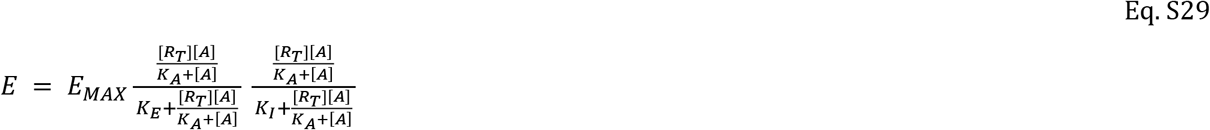

That simplifies to Eq. S30.

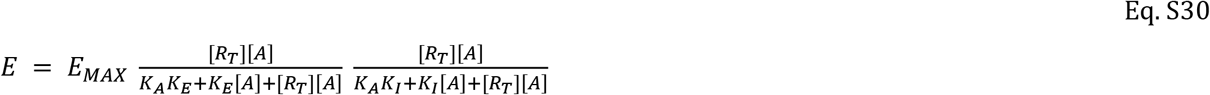

Division of Eq. S30 by K_E_ and K_I_ and substitution τ = [R_T_]/K_E_ and σ = [R_T_]/K_I_ gives Eq. S31.

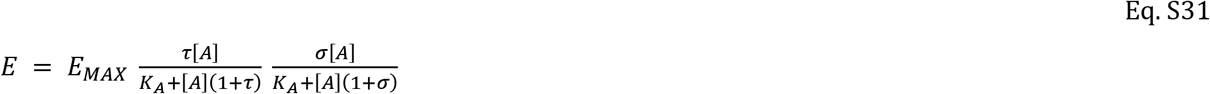

Apparent E’_MAX_ at [A]>>K_A_ is given by Eq. S32.

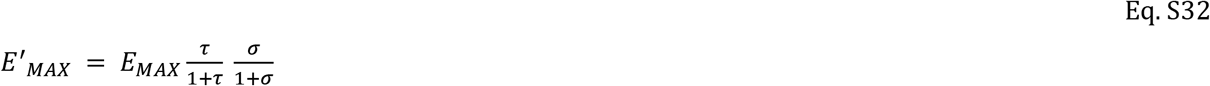

The apparent value of operational efficacy τ’ is given by equation Eq. S33.

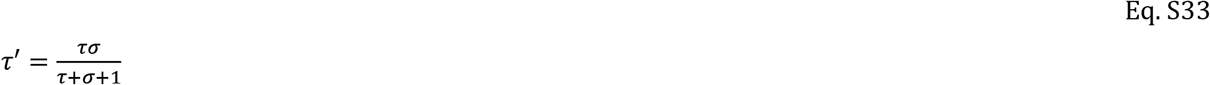

For half-efficient concentration EC_50_:

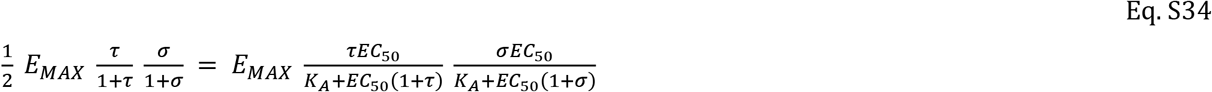

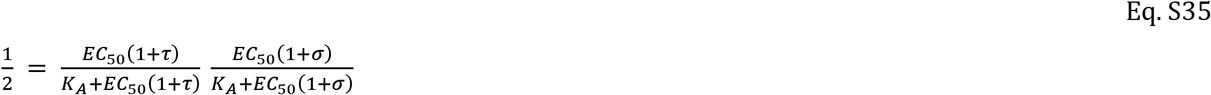

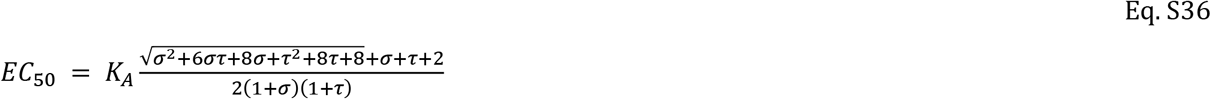

**Figure S7.**
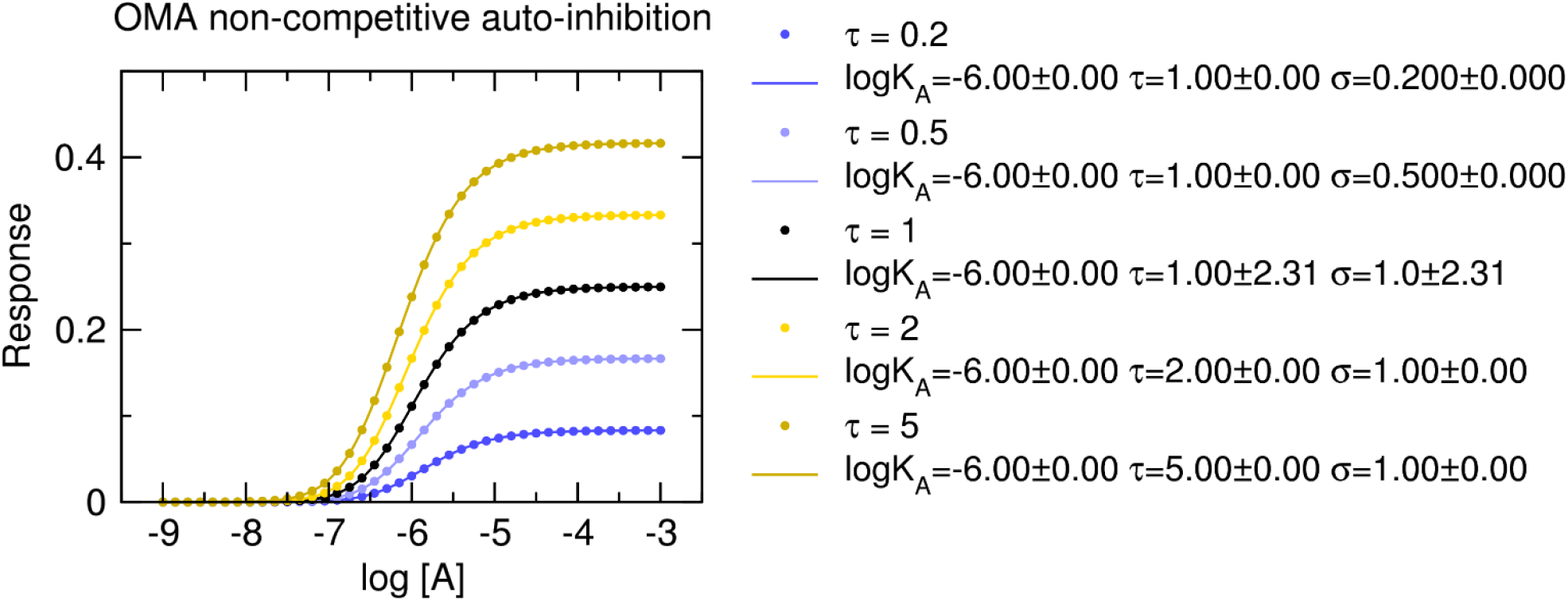
Fitting Eq. S31 to the model of non-competitive auto-inhibition. Dots, functional-response data modelled according to Eq. S29. E_MAX_ = 1, K_A_ = 10^-6^M, σ = 1, R_T_ =1. Values of operational efficacy τ are indicated in the legend. Lines, fits of Eq. S31 to the model data. E_MAX_ was fixed to 1, initial estimate of τ was set to 0.3. Parameter estimates of the fits are indicated in the legend.

Parameter estimates are correct, associated with low level of uncertainty. However, it is result of initial estimate of τ. For K_I_=5, estimates of operational efficacy τ and inhibition factor σ are swapped pointing to the symmetry of Eq. S31.

**Figure S8.**
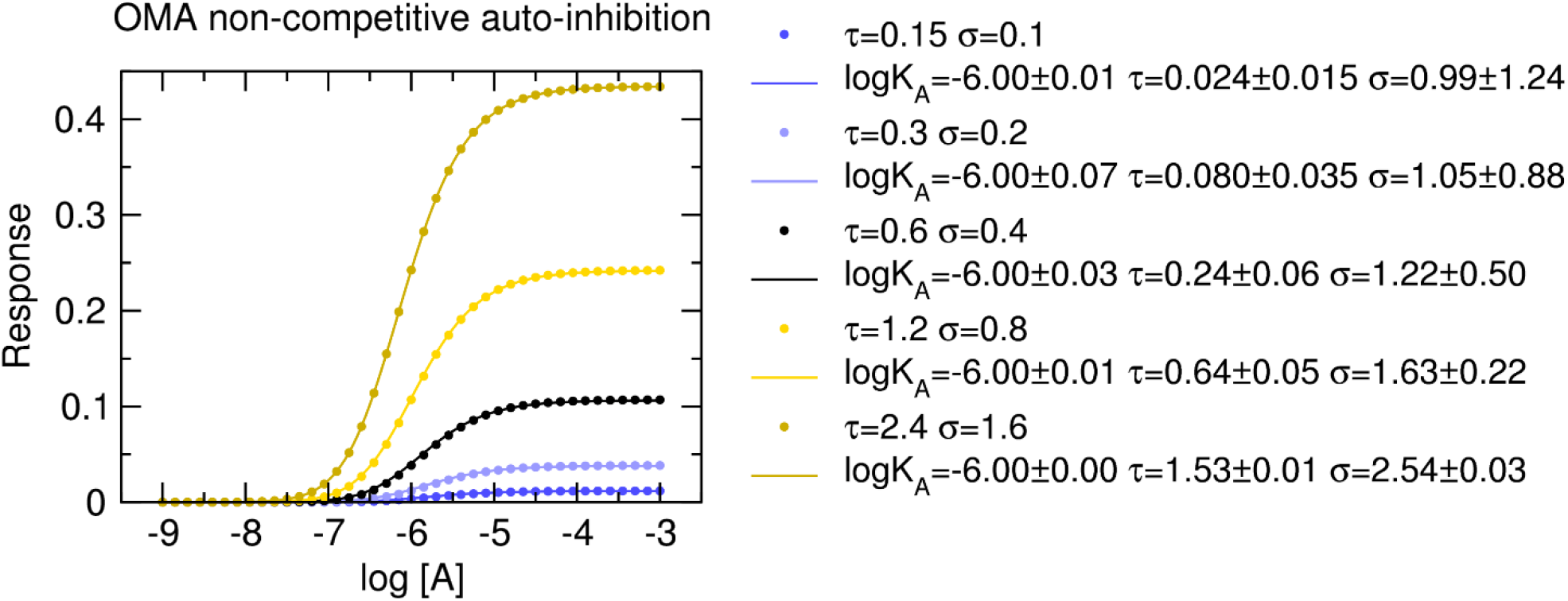
Fitting Eq. S31 to the model of non-competitive auto-inhibition with varying receptor concentration. Dots, functional-response data modelled according to Eq. S29. E_MAX_ = 1, K_A_ = 10^-6^M, K_E_ = 3.333, K_I_ = 5. R_T_ varied from 0.5 to 8. Resulting operational efficacies τ and inhibition factors σ are indicated in the legend. Lines, fits of Eq. S31 to the model data. E_MAX_ was fixed to 1, initial estimate of τ was set to apparent operational efficacy τ’ calculated as E’_MAX_/(E_MAX_-E’_MAX_). Parameter estimates of the fits are indicated in the legend.

Parameter estimates are incorrect and for low values of τ and σ they are associated with high level uncertainty. However, calculated τ and σ values give correct apparent efficacy according to Eq. S33.

### Signalling feedback

In signalling feedback, an increase in output signal proportionally either decreases (negative feedback) or increases (positive feedback) input. Activation constant K_E_ can be expressed as difference between [RAE] formation and decay Eq. S 37.

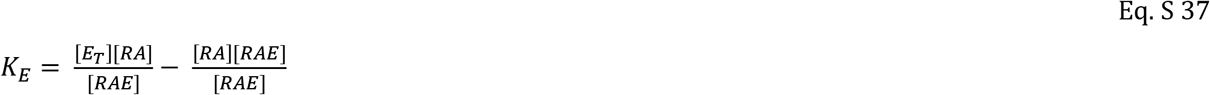

Where [E_T_] is total concentration of effector. Eq. S 37can be simplified to Eq. S 38.

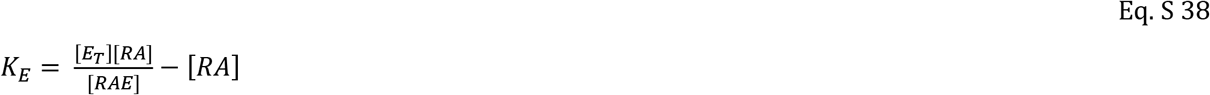

After rearrangement

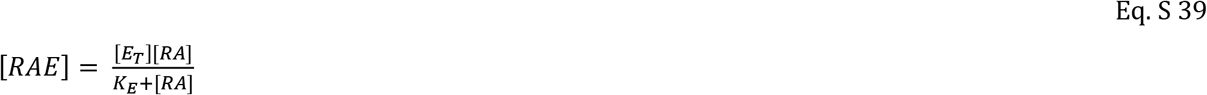

The feedback factor δ modifies input [RA]

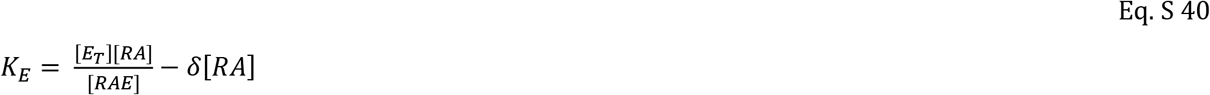

Respectively

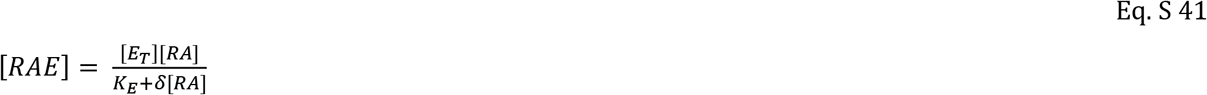

For δ > 1, right member of Eq. S 40 is greater than right member of Eq. S 38 and thus denotes attenuation of the signal, negative feedback. Conversely, δ < 1 denotes positive feedback. The proportion of signal output, [RAE], working as feedback is constant and is given by division of Eq. S 41 by Eq. S 39.

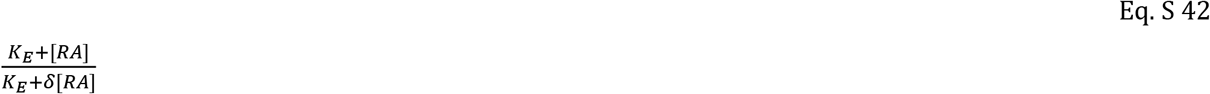

Under the feedback, response is given by:

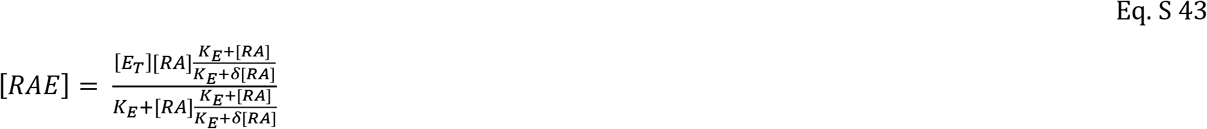

After simplification

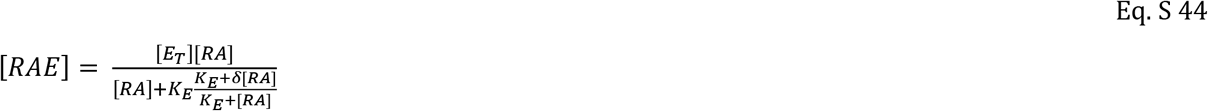

Taking [RAE] as a functional response E and [E_T_] as maximal response of the system E_MAX_ and substitution binding equation (Eq. S5) into Eq. S 44 we receive:

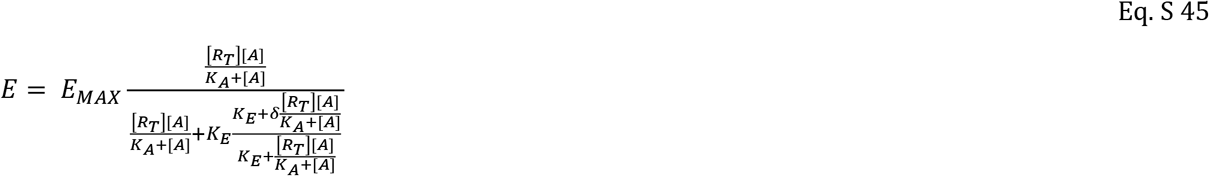

After simplification:

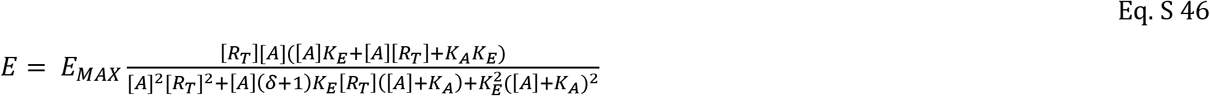

After multiplication of parentheses:

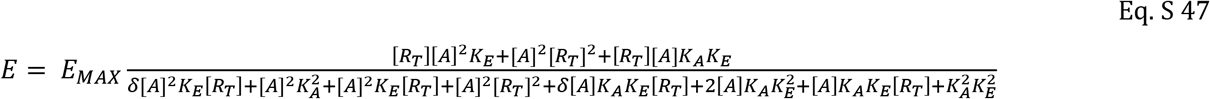

After division by K_E_^2^ and substitution τ = [R_T_]/K_E_:

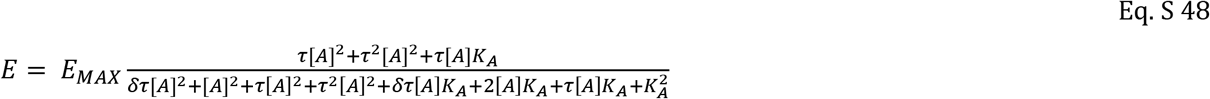

After simplification:

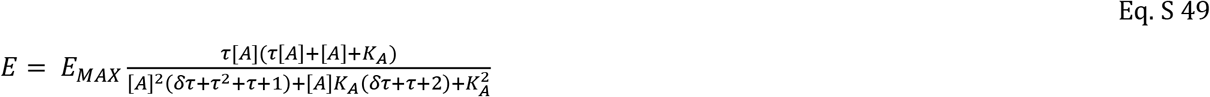

For [A]>>K_A_ Eq. S 49 becomes:

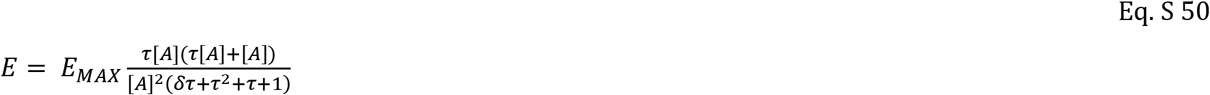

Thus, maximal response to an agonist with operational efficacy τ is:

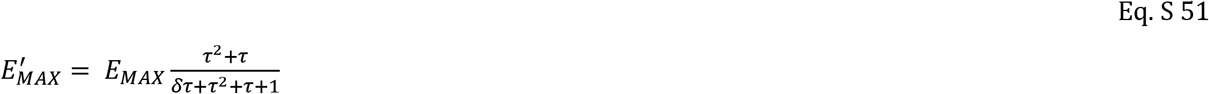

And apparent value of operational efficacy is:

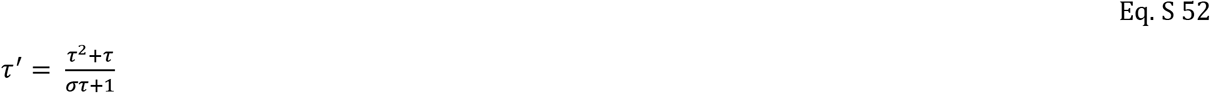

Solving Eq. S 49 for ½ E’_MAX_ gives EC_50_ value as:

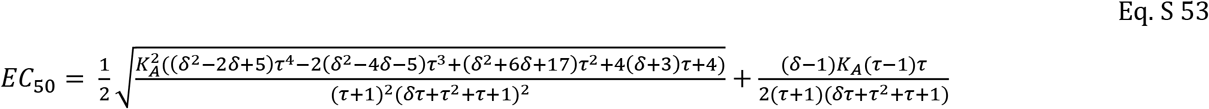

Alternatively,

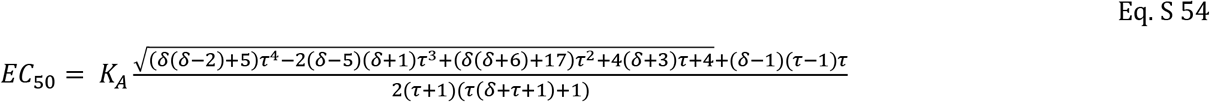

**Figure S9.**
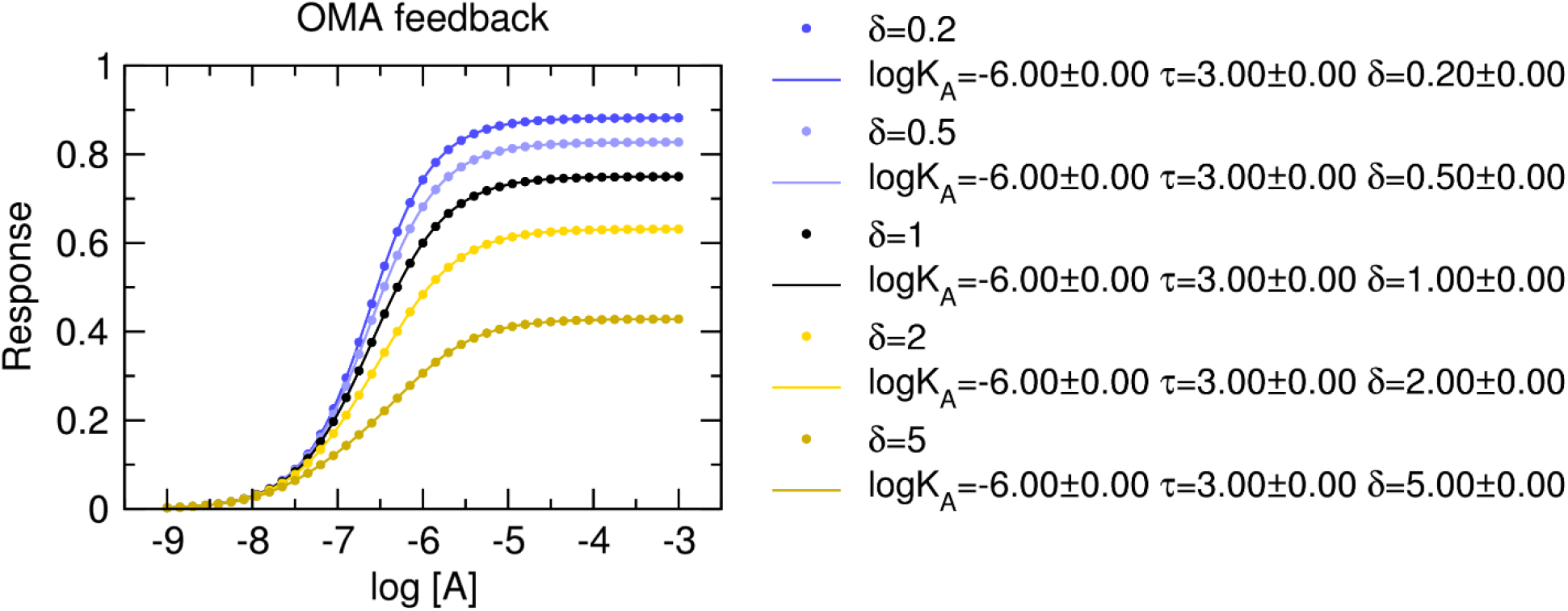
Fitting Eq. S 49 to the signalling-feedback model with varying feedback. Dots, functional-response data modelled according to Eq. S 45. E_MAX_ = 1, K_A_ = 10^-6^M, K_E_ = 3.333, R_T_ = 1. Feedback factor δ varied from 0.2 to 5 and is indicated in the legend. Lines, fits of Eq. S 49 to the model data. E_MAX_ was fixed to 1, initial estimate of τ was set to 3 and initial estimate of logK_A_ was set to −6. Parameter estimates of the fits are indicated in the legend.

**Figure S10.**
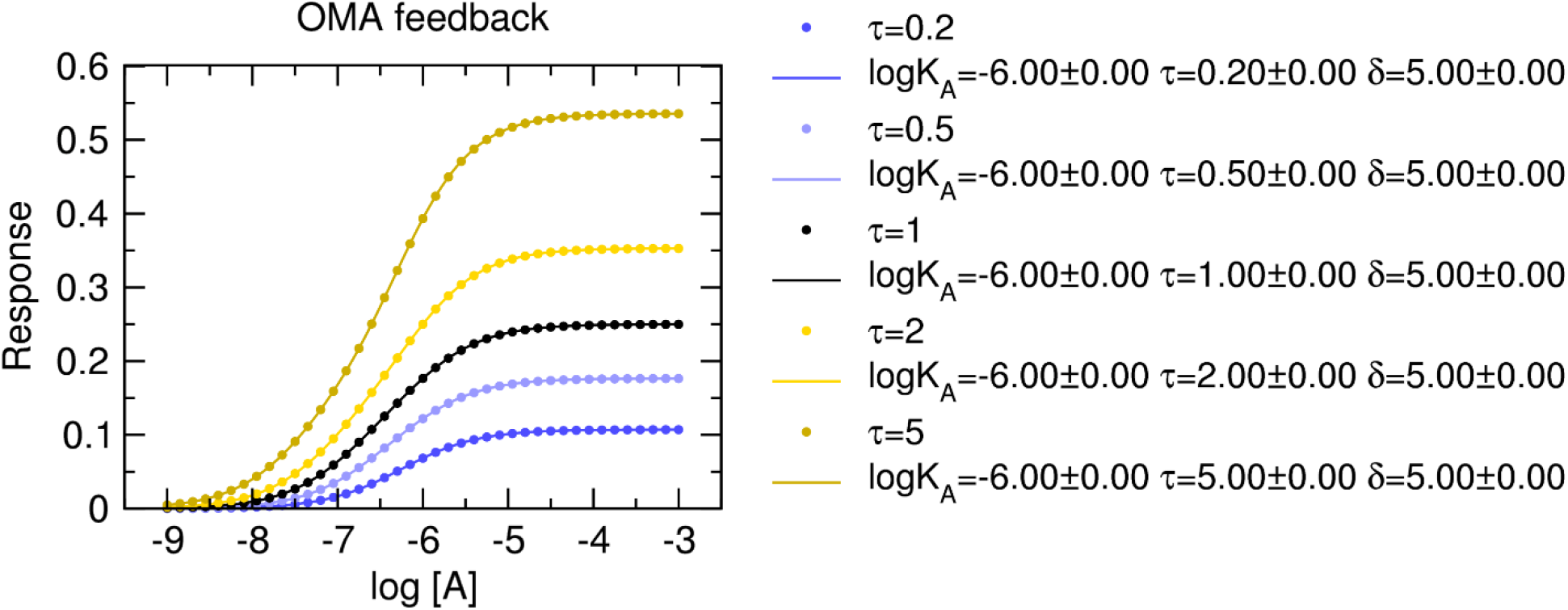
Fitting Eq. S 49 to the signalling-feedback model with constant feedback and varying operational efficacy. Dots, functional-response data modelled according to Eq. S 45. E_MAX_ = 1, K_A_ = 10^-6^M, R_T_ = 1, δ=5. Operational efficacy varied from 0.2 to 5 and is indicated in the legend. Lines, fits of Eq. S 49 to the model data. E_MAX_ was fixed to 1, initial estimate of δ was set to 5 and initial estimate of logK_A_ was set to −6. Parameter estimates of the fits are indicated in the legend.

### Low receptor-expression systems

Equation Eq. S6 is valid only when [RA]>>[E]. For [RA]≈[E] K_E_ is given by:

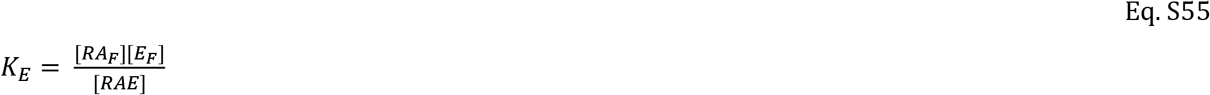

Where [RA_F_] and [E_F_] are free concentrations of RA and E, respectively, and are given by:

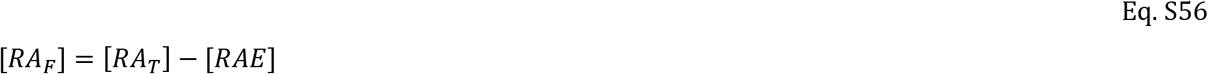

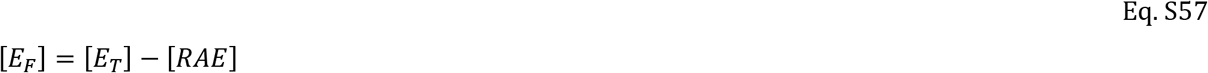

Where [RA_T_] and [E_T_] are total concentrations of RA and E, respectively, and RAE is complex of RA and E. By substitution Eq. S56 and Eq. S57 into Eq. S55:

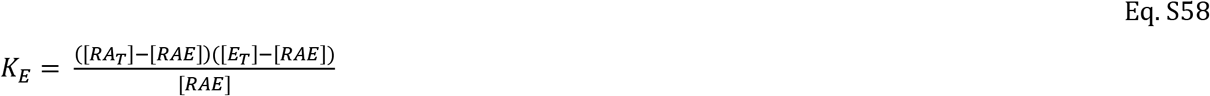

Rearranging:

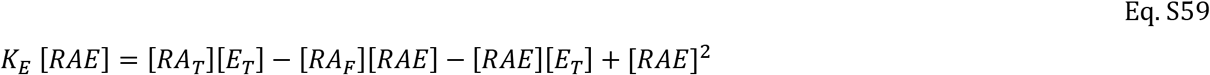

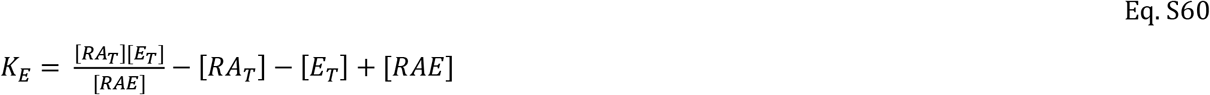

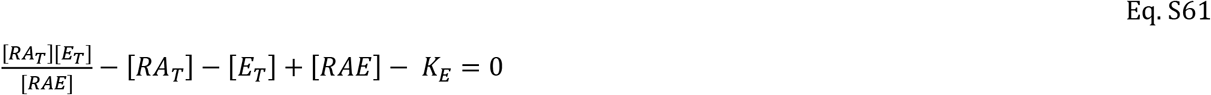

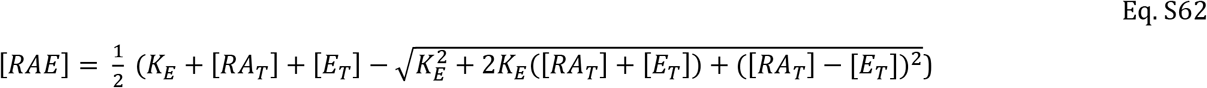

Substituting binding function (Eq. S5) for [RA_T_] into Eq. S61 gives:

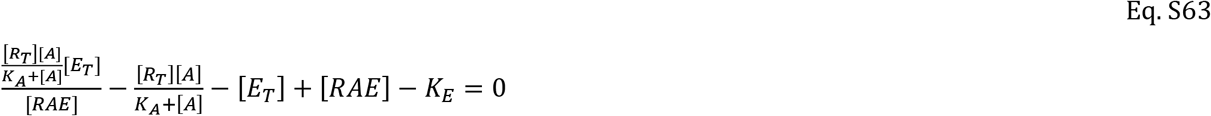

That simplifies to:

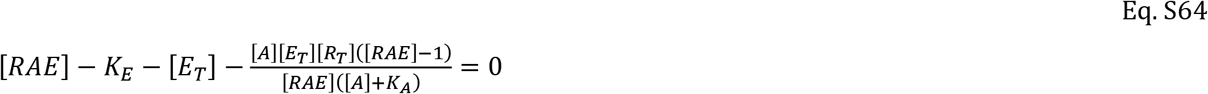

Solving Eq. S64 for [RAE] gives only approximate solutions.

**Figure S 11.**
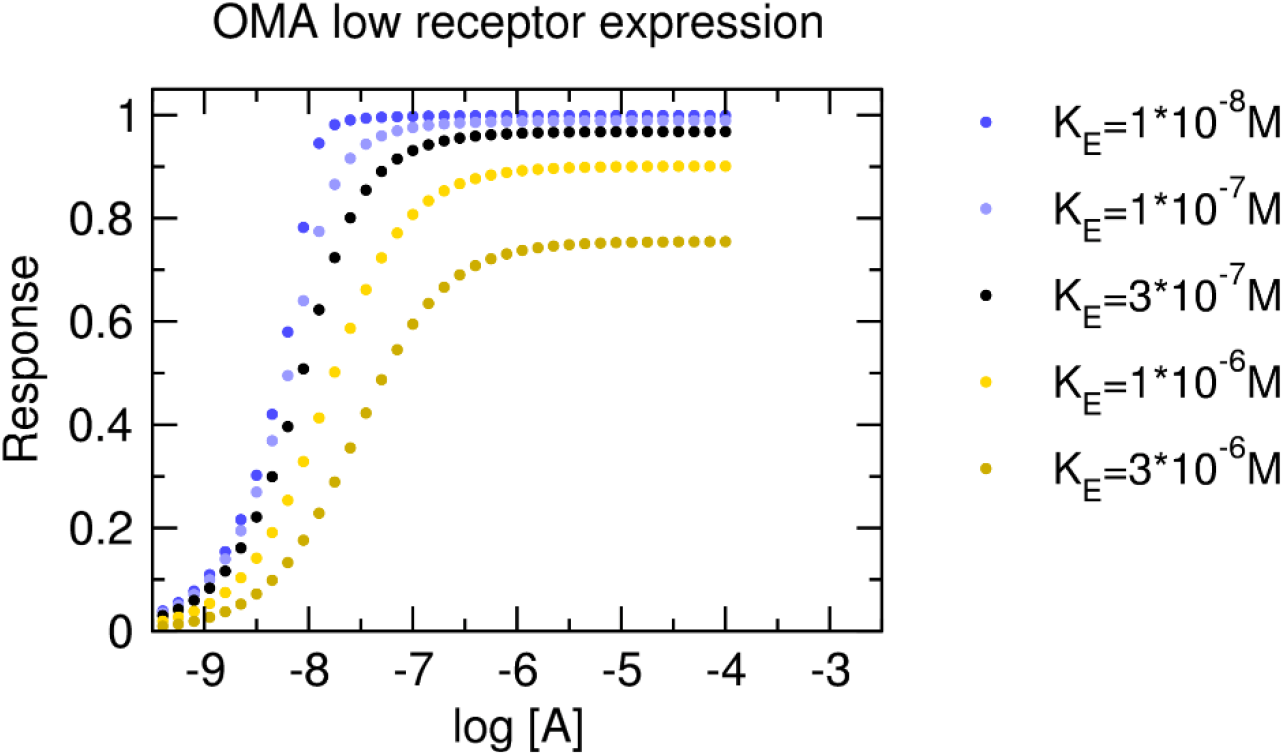
Modelling the system with a low expression of receptors. Dots, functional-response data modelled in two steps. First, binding was calculated according to Eq. S5. Then resulting [RA] was used in Eq. S62. E_T_ = 10^-6^M, R_T_ = 10^-5^M, K_A_ = 10^-6^M. Values of K_E_ are indicated in the legend. As Eq. S64 gives only approximate solution, data were not refitted.

### Python scripts

Scripts to model functional-response data and fit equations of Black & Leff, Hill and explicit models to them are provided in python.zip.

Requirements: python, numpy, matplotlib, scipy

Content of python.zip:

Folders:

- Black-Leff
- Hill
- OMA_low-expression_FR
- OMA_low-expression_Ke
- OMA_NCI
- OMA_Rtot
- OMA_SInh

#### Black-Leff

Black-Leff_generate_5_data_sets.py generates 5 datasets named Data_A.dat through Data_E.dat according to the Black & Leff operational model of agonism (OMA) with parameters in the header of the script. Parameters of the model are saved in Black-Leff.par.

#### Hill

Hill_generate_5_data_sets.py generates 5 datasets named Data_A.dat through Data_E.dat according to the OMA modified with the Hill coefficient with parameters in the header of the script. Parameters of the model are saved in Hill.par.

#### OMA_low-expression_FR

OMA_low-expression_FR_generate_5_data_sets.py generates 5 datasets of functional response named Data_A.dat through Data_E.dat according to the OMA of system expressing receptors in low concentration with parameters in the header of the script. Parameters of the model are saved in OMA_low-expression_FR.par.

Black-Leff_fit.py fits the Black & Leff OMA to the data. Fit results are saved in Black-Leff_fit.res Fit curves are saved as *_Black-Leff_fit.data.

Hill_fit.py fits the OMA modified with the Hill coefficient to the data. Fit results are saved in Hill_fit.res Fit curves are saved as *_Hill_fit.data.

#### OMA_low-expression_Ke

OMA_low-expression_Ke_generate_5_data_sets.py generates 5 datasets of signal transduction named Data_A.dat through Data_E.dat according to the OMA of system expressing receptors in low concentration with parameters in the header of the script. Parameters of the model are saved in OMA_low-expression_Ke.par.

Black-Leff_fit.py fits the Black & Leff OMA to the data. Fit results are saved in Black-Leff_fit.res Fit curves are saved as *_Black-Leff_fit.data.

Hill_fit.py fits the OMA modified with the Hill coefficient to the data. Fit results are saved in Hill_fit.res Fit curves are saved as *_Hill_fit.data.

#### OMA_NCI

OMA_NCI_generate_5_data_sets.py generates 5 datasets named Data_A.dat through Data_E.dat according to the OMA of non-competitive auto-inhibition with parameters in the header of the script. Parameters of the model are saved in OMA_NCI.par.

Black-Leff_fit.py fits the Black & Leff OMA to the data. Fit results are saved in Black-Leff_fit.res Fit curves are saved as *_Black-Leff_fit.data.

Hill_fit.py fits the OMA modified with the Hill coefficient to the data. Fit results are saved in Hill_fit.res Fit curves are saved as *_Hill_fit.data.

OMA_NCI_fit.py fits the OMA of non-competitive auto-inhibition to the data. Fit results are saved in OMA_NCI_fit.res Fit curves are saved as *_OMA_NCI_fit.data.

#### OMA_NCI_Rtot

OMA_NCI_Rtot_generate_5_data_sets.py generates 5 datasets named Data_A.dat through Data_E.dat according to the OMA of non-competitive auto-inhibition with varying number of receptors Rtot with parameters in the header of the script. Parameters of the model are saved in OMA_NCI.par.

Black-Leff_fit.py fits the Black & Leff OMA to the data. Fit results are saved in Black-Leff_fit.res Fit curves are saved as *_Black-Leff_fit.data.

Hill_fit.py fits the OMA modified with the Hill coefficient to the data. Fit results are saved in Hill_fit.res Fit curves are saved as *_Hill_fit.data.

OMA_NCI_Rtot_fit.py fits the OMA of non-competitive auto-inhibition with varying Rtot to the data. Fit results are saved in OMA_NCI_Rtot_fit.res Fit curves are saved as *_OMA_NCI_Rtot_fit.data.

#### OMA_SInh

OMA_SInh_generate_5_data_sets.py generates 5 datasets named Data_A.dat through Data_E.dat according to the OMA of substrate inhibition with parameters in the header of the script. Parameters of the model are saved in OMA_SInh.par.

Black-Leff_fit.py fits the Black & Leff OMA to the data. Fit results are saved in Black-Leff_fit.res Fit curves are saved as *_Black-Leff_fit.data.

Hill_fit.py fits the OMA modified with the Hill coefficient to the data. Fit results are saved in Hill_fit.res Fit curves are saved as *_Hill_fit.data.

OMA_SInh_fit.py fits the OMA of substrate inhibition to the data. Fit results are saved in OMA_SInh_fit.res Fit curves are saved as *_OMA_SInh_fit.data.

## Notes

### Competing Interest Statement

The authors have declared no competing interest.

### Summary of Updates

The section on allosteric modulation was removed. Sections on signalling feedback and low receptor-expression systems were added. The discussion was rewritten.

## References

Black, J.W., and Leff, P. (1983). Operational models of pharmacological agonism. Proc. R. Soc. London. Ser. B, Biol. Sci. 220: 141–62.

Black, J.W., Leff, P., Shankley, N.P., and Wood, J. (1985). An operational model of pharmacological agonism: the effect of E/[A] curve shape on agonist dissociation constant estimation. Br. J. Pharmacol. 84: 561–71.

Burgueño, J., Pujol, M., Monroy, X., Roche, D., Varela, M.J., Merlos, M., et al. (2017). A Complementary Scale of Biased Agonism for Agonists with Differing Maximal Responses. Sci. Rep. 7: 15389.

Christopoulos, A., and El-Fakahany, E.E. (1999). Qualitative and quantitative assessment of relative agonist efficacy. Biochem. Pharmacol. 58: 735–748.

Ehlert, F.J., Griffin, M.T., Sawyer, G.W., and Bailon, R. (1999). A simple method for estimation of agonist activity at receptor subtypes: comparison of native and cloned M3 muscarinic receptors in guinea pig ileum and transfected cells. J. Pharmacol. Exp. Ther. 289: 981–92.

Furchgott, R.F. (1966). The use of β-haloalkylamines in the diferentiation of receptors and in the determination of dissociation constants of receptor-agonist complexes. Adv. Drug Res. 3: 21–55.

Gesztelyi, R., Zsuga, J., Kemeny-Beke, A., Varga, B., Juhasz, B., and Tosaki, A. (2012). The Hill equation and the origin of quantitative pharmacology. Arch. Hist. Exact Sci. 66: 427–438.

Gregory, K.J., Giraldo, J., Diao, J., Christopoulos, A., and Leach, K. (2020). Evaluation of Operational Models of Agonism and Allosterism at Receptors with Multiple Orthosteric Binding Sites. Mol. Pharmacol. 97: 35–45.

Griffin, M.T., Figueroa, K.W., Liller, S., and Ehlert, F.J. (2007). Estimation of agonist activity at G protein-coupled receptors: analysis of M2 muscarinic receptor signaling through Gi/o,Gs, and G15. J. Pharmacol. Exp. Ther. 321: 1193–207.

Jakubík, J., and Randáková, A. (2022). Insights into the operational model of agonism of receptor dimers. Expert Opin. Drug Discov. 17: 1181–1191.

Jakubík, J., Randáková, A., Chetverikov, N., El-Fakahany, E.E., and Doležal, V. (2020). The operational model of allosteric modulation of pharmacological agonism. Sci. Rep. 10: 14421.

Jakubík, J., Randáková, A., Rudajev, V., Zimčík, P., El-Fakahany, E.E., and Doležal, V. (2019). Applications and limitations of fitting of the operational model to determine relative efficacies of agonists. Sci. Rep. 9: 4637.

Kenakin, T. (2017). A Scale of Agonism and Allosteric Modulation for Assessment of Selectivity, Bias, and Receptor Mutation. Mol. Pharmacol. 92: 414–424.

Kenakin, T., and Christopoulos, A. (2013). Signalling bias in new drug discovery: detection, quantification and therapeutic impact. Nat. Rev. Drug Discov. 12: 205–16.

Kenakin, T., Watson, C., Muniz-Medina, V., Christopoulos, A., and Novick, S. (2012). A simple method for quantifying functional selectivity and agonist bias. ACS Chem. Neurosci. 3: 193–203.

Kenakin, T.P. (2012). Biased signalling and allosteric machines: New vistas and challenges for drug discovery. Br. J. Pharmacol. 165: 1659–1669.

Kenakin, T.P. (2014). Agonists: The Measurement of Affinity and Efficacy in Functional Assays. In A Pharmacology Primer, (Academic Press), pp 85–117.

Keov, P., López, L., Devine, S.M., Valant, C., Lane, J.R., Scammells, P.J., et al. (2014). Molecular mechanisms of bitopic ligand engagement with the M1 muscarinic acetylcholine receptor. J. Biol. Chem. 289: 23817–37.

Lefkowitz, R.J. (2013). A brief history of G-protein coupled receptors (Nobel Lecture). Angew. Chem. Int. Ed. Engl. 52: 6366–78.

Luttrell, L.M., Maudsley, S., and Bohn, L.M. (2015). Fulfilling the Promise of ‘Biased’ G Protein-Coupled Receptor Agonism. Mol. Pharmacol. 88: 579–88.

Roche, D., Graaf, P.H. van der, and Giraldo, J. (2016). Have many estimates of efficacy and affinity been misled? Revisiting the operational model of agonism. Drug Discov. Today 21: 1735–1739.

Stott, L.A., Hall, D.A., and Holliday, N.D. (2016). Unravelling intrinsic efficacy and ligand bias at G protein coupled receptors: A practical guide to assessing functional data. Biochem. Pharmacol. 101: 1–12.

